# SETDB1 activity is globally directed by H3K14 acetylation via its Triple Tudor Domain

**DOI:** 10.1101/2024.04.22.590554

**Authors:** Thyagarajan T. Chandrasekaran, Michel Choudalakis, Alexander Bröhm, Sara Weirich, Alexandra G Kouroukli, Ole Ammerpohl, Philipp Rathert, Pavel Bashtrykov, Albert Jeltsch

**Affiliations:** Institute of Biochemistry and Technical Biochemistry, University of Stuttgart, Allmandring 31, 70569 Stuttgart, Germany; Institute of Human Genetics, University of Ulm and Ulm University Medical Center, Albert-Einstein-Allee 11, 89091 Ulm, Germany

**Keywords:** Protein lysine methyltransferase, Histone methylation, H3K9me3, H3K14ac, repeat elements

## Abstract

SETDB1 is a major H3K9 methyltransferase involved in heterochromatin formation and silencing of repeat elements. It contains a unique Triple Tudor Domain (3TD) which specifically binds the dual modification of H3K14ac in the presence of H3K9me1/2/3. Here, we explored the role of the 3TD H3-tail interaction for the H3K9 methylation activity of SETDB1. We generated a binding reduced 3TD mutant and demonstrate in biochemical methylation assays on peptides and recombinant nucleosomes containing H3K14ac analogs, that H3K14 acetylation is crucial for the 3TD mediated recruitment of SETDB1. We also observe this effect in cells where SETDB1 binding and activity is globally correlated with H3K14ac, and KO of the H3K14 acetyltransferase HBO1 causes a drastic reduction in H3K9me3 levels at SETDB1 dependent sites. Further analyses revealed that 3TD particularly important at specific target regions like L1M repeat elements, where SETDB1 KO cannot be efficiently reconstituted by the 3TD mutant of SETDB1. In summary, our data demonstrate that the H3K9me3 and H3K14ac are not antagonistic marks but rather the presence of H3K14ac is required for SETDB1 recruitment via 3TD binding to H3K9me1/2/3-K14ac and establishment of H3K9me3.

## Introduction

Histone tails are pivotal in regulation of gene expression through a variety of post-translational modifications (PTMs), including acetylation, methylation, and phosphorylation ^1,2^. These modifications, which occur on specific amino acids such as lysine, arginine, serine, or threonine, have a variety of essential roles in development and disease ^3,4^. Among the enzymes modifying histones, the SETDB1 (SET domain bifurcated histone lysine methyltransferase 1) protein lysine methyltransferase (PKMT) stands out for its role in trimethylating lysine 9 on histone H3 (H3K9me3) ^5,6^. H3K9me3 is a critical heterochromatic histone PTM crucial for silencing of repeat elements (RE), embryonic development and its dysregulation is frequently associated with cancer proliferation ^7–9^. Consequently, SETDB1 has frequently been associated with these processes ^5,10–14^. The recruitment of SETDB1 to specific genomic sites involves diverse mechanisms, including interaction with the HUSH complex for repression of LINE1 elements ^6,15,16^ and intron-less genes ^17^. In addition, SETDB1 is recruited to endogenous retroviruses (ERVs) by KAP1 (also known as TRIM28) through binding of KRAB-ZFP proteins ^18–20^. Additional recruitment of epigenetic regulators to the SETDB1 complex such as DNMT3A and the NuRD deacetylase complex ^21,22^ results in formation of a comprehensive repressive complex with multiple functionality.

SETDB1 comprises a unique Triple Tudor Domain (3TD), a methyl-CpG-binding domain (MBD) and a SET domain (Figure 1A) ^5,6^. The SET domain of SETDB1 harboring its catalytic activity is split into SET-N and SET-C parts separated by a large insert region, SET-I, which together contribute to target lysine recognition and methylation. While the function of the MBD domain has not been shown so far, 3TD has been found to specifically recognize the dual modification consisting of H3K14ac in the presence of H3K9me1/2/3 ^23^. H3K14ac provides a primary signal for 3TD binding in this dual mark, because binding of peptides only containing H3K14ac was observed, while peptides only containing H3K9me1/2/3 were not bound ^23^. Crystal structures of 3TD in complex with H3K14ac-K9me2/3 peptides demonstrated how peptide binding and K14ac recognition specifically occurs at the interface between the second and third Tudor domains of 3TD by forming a direct hydrophobic interaction with F332 (Supplementary Figure 1A). Indeed, an F332A exchange led to a strong reduction in binding affinity of 3TD to modified H3 peptides ^23^.

**Figure 1:**
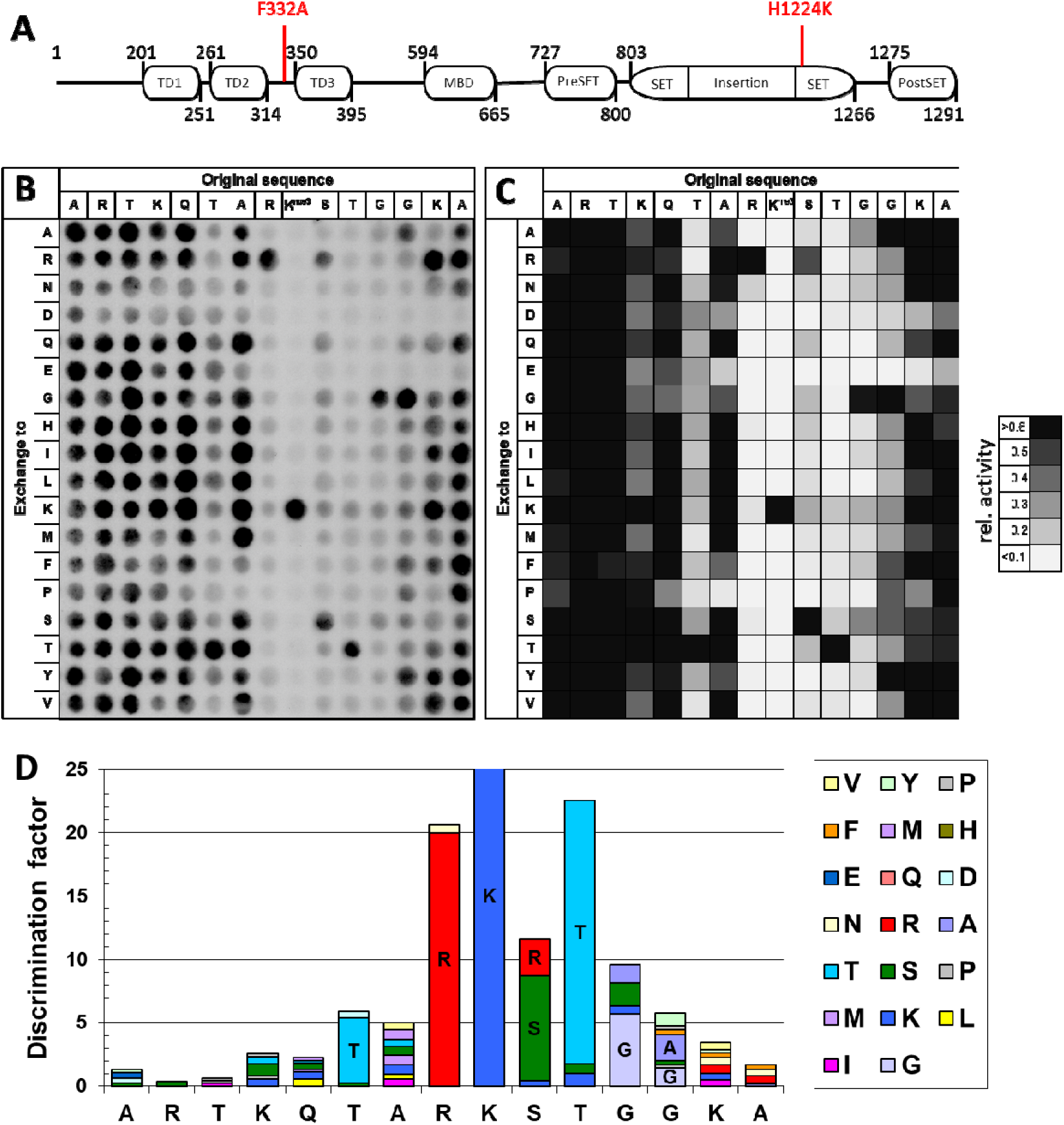
Specificity analysis of human SETDB1. **A)** Schematic representation of SETDB1 domains. The mutants used in this study are highlighted in red (based on Uniprot Q15047 and ^23^). **B)** Substrate sequence specificity analysis of full-length SETDB1 (see also Supplementary Figure 1B) using SPOT peptide array methylation of H3 (1-20) peptides. The array contains a systematic collection of single mutants of position 1-15 of target sequence in which each amino acid residue was exchanged against 18 natural amino acids. The peptide array was methylated by SETDB1 in methylation buffer containing radioactively labelled AdoMet and the transfer of methyl group was detected by autoradiography. **C)** Analysis of two independent peptide array methylation experiments, where the signals were quantified, averaged, normalized and indicated in greyscale (see also Supplementary Figure 1C). **D)** Discrimination factor analysis showing the sequence substrate specificity of SETDB1 in methylating the H3K9 peptide.

The H3K9me1/2/3-K14ac dual modification is a highly abundant modification in the human genome ^24–26^ adding potential biological relevance to the biochemical identification of the 3TD dual specificity. Interestingly, the H3K9me1/2/3-K14ac binding preference of 3TD include H3K9me2/K14ac and H3K9me3/K14ac, which both represent highly abundant bivalent modifications. The term bivalent is used for apparently opposing histone marks occurring together at one genomic locus ^27,28^. It has been reported in mice that the bivalent modification of H3K9me3 and H3K14ac marks the poised inactive state of the genome ^29^.

In this study, we explored the role of the 3TD H3K9me1/2/3-K14ac interaction in the H3K9 methylation activity of SETDB1. We utilized the 3TD binding reduced F332A mutant and studied the interplay of 3TD and SET domain of SETDB1 in substrate recognition and methylation. Our data reveal that H3K14 acetylation is crucial for the recruitment of SETDB1 to its targets and to efficiently introduce H3K9 methylation, as evidenced our by biochemical methylation assays using recombinant nucleosomes containing H3K14ac analogs. We also observed this effect in cellular assays with SETDB1 KO cells lines where H3K14ac was globally correlated with SETDB1 activity and binding. Strikingly, KO of the main H3K14 acetyltransferase HBO1 (also known as KAT7) caused a drastic reduction in the H3K9me3 levels at SETDB1 dependent sites. Further analysis revealed that 3TD was not required for SETDB1 recruitment to genomic regions targeted by KAP1. However, at specific groups of target regions, SETDB1 KO could not be efficiently reconstituted by SETDB1 mutants containing a the F332A mutation in 3TD indicating that the intact chromatin interaction of 3TD is essential for SETDB1 recruitment. Our data show that H3K9 methylation at L1M RE is highly dependent on an intact 3TD. In summary, we demonstrate an additional mechanism of SETDB1 recruitment to chromatin through the 3TD interaction with H3 tails containing K14ac and K9 methylation which is particularly relevant at L1M RE.

## Results and discussion

### H3K9 methylation specificity profile of SETDB1

The catalytic domains of PKMTs interact with the amino acid residues surrounding the lysine methylation site of substrate proteins or peptides and this sequence specific interaction is a critical parameter to define the substrate specificity of PKMTs ^30^. However, the exact preferences for certain amino acid residues at each position of the methylation substrate are not well studied for many PKMTs including SETDB1. We, therefore, wanted to study the substrate sequence specificity of full-length SETDB1 which was expressed in insect cells and purified by affinity chromatography (Supplementary Figure 1B). Peptide SPOT arrays were employed as powerful approach to map the substrate specificity of PKMTs ^31,32^. In this experiment, the natural H3 (1-20 aa) was used as a template sequence and in each spot one single amino acid in the 1-15 region was exchanged to 18 other natural amino acids, creating all possible single amino acid mutants of the 1-15 region of the template sequence. Final SPOT arrays contained 270 (15 x 18) different peptides whose methylation was studied in a competitive manner. Methylation was detected by incubating the SPOT arrays with purified SETDB1 in the methylation buffer containing radioactively labelled AdoMet as a cofactor and the transfer of methyl groups was detected by autoradiography (Figure 1B). The methylation experiments were performed in duplicates which were then normalized and averaged (Figure 1C). The error distribution was calculated for each spot showing that most spots had SD smaller than ± 10% (Supplementary Figure 1C) indicating high reproducibility of the results. Finally, a discrimination factor ^31^ was calculated illustrating the enzyme specificity in form of the preference of SETDB1 for individual amino acids in each position of H3 peptide over all the other amino acids at the corresponding position (Figure 1D).

As expected, an exchange of the target lysine K9 by any other amino acid led to a complete loss of enzymatic activity on the corresponding peptides (Figure 1B and C). Like the other H3K9 PKMTs SUV39H1 ^33^, SUV39H2 ^34^, and G9a ^35^, SETDB1 showed a high specificity for arginine at the -1 position (the target K9 is considered as the reference position). Similar to SUV39H2, SETDB1 shows additional sequence readout from +1 to +4 position, with a preference for S at the +1, strong preference for T at +2, preference for G at +3 and for G/A at +4. Moreover, SETDB1 demonstrated a unique preference for T at the -3 position. At some of these sites, alternative residues were accepted, though with reduced efficiency, yielding the following recognition motif:

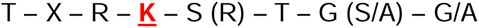

Hence, the SET domain of SETDB1 shows highly specific sequence readout from the -3 to +4 position of the H3 peptide. Of note, there is no sequence readout by the SET domain at the K14 position. In addition, as explained above SETDB1 contains a 3TD reader domain that contributes to the target recognition ^23^. As shown in previous biochemical and structural data, the 3TD of SETDB1 specifically binds to the H3 peptide containing K14ac-K9me1/2/3 and it interacts with amino acids ranging from the K9 to the +8 position ^23^. This indicates that the SET and 3TD domains of SETDB1 have an overlapping substrate binding profile strongly suggesting a competition of both domains for peptide interaction if the sequence contains K14ac.

### Peptide and recombinant nucleosome methylation by SETDB1

Next, we wanted to understand the role of H3K9me1/2/3-K14ac binding by 3TD in the methylation of the target H3K9. For this, we used the SETDB1 3TD mutant F332A which disrupts a contact of 3TD to the double modified H3 peptide (Supplementary Figure 1A) and at domain level showed a strong reduction in binding to peptides containing H3K9me1/2/3-K14ac ^23^. Full-length SETDB1 F332A was generated, expressed in insect cells, purified and used in biochemical activity assays at same concentrations as WT SETDB1 (Supplementary Figure 1B). Unmodified H3 peptides (1-18) along with K14ac modified H3 (1-18) peptides were incubated with SETDB1 WT or F332A in the methylation buffer containing radioactively labelled AdoMet as a cofactor and transfer of methyl groups were then detected by autoradiography. Peptides containing either only K9me3 or both K9me3 and K14ac were used as negative controls.

The peptide methylation data shown in Figure 2A clearly indicate that SETDB1 WT methylates the unmodified H3 peptide. As expected, no methylation signals were observed with peptides containing H3K9me3 confirming the specificity of SETDB1 towards methylation of H3K9. Interestingly, we observed a drastic reduction in the methylation signal of the peptide containing H3K14ac in case of SETDB1 WT (Figure 2A), which agrees with previous data ^21^. In contrast, only a moderate reduction of the methylation signal was observed with the 3TD mutant F332A (Figure 2A). The competition of the 3TD and SET domains for binding to the H3K14ac containing H3 peptides can explain the differential methylation activity of SETDB1 WT and F332A on H3 peptides with or without H3K14ac, because in the case of WT SETDB1, the presence of H3K14ac leads to 3TD binding which then hinders the access of the SET domain to the substrate causing an inhibition of the catalytic activity. In contrast, in the case of the 3TD mutant F332A, due to the strong reduction of binding to modified H3 peptides, the 3TD mediated inhibition is much weaker.

**Figure 2:**
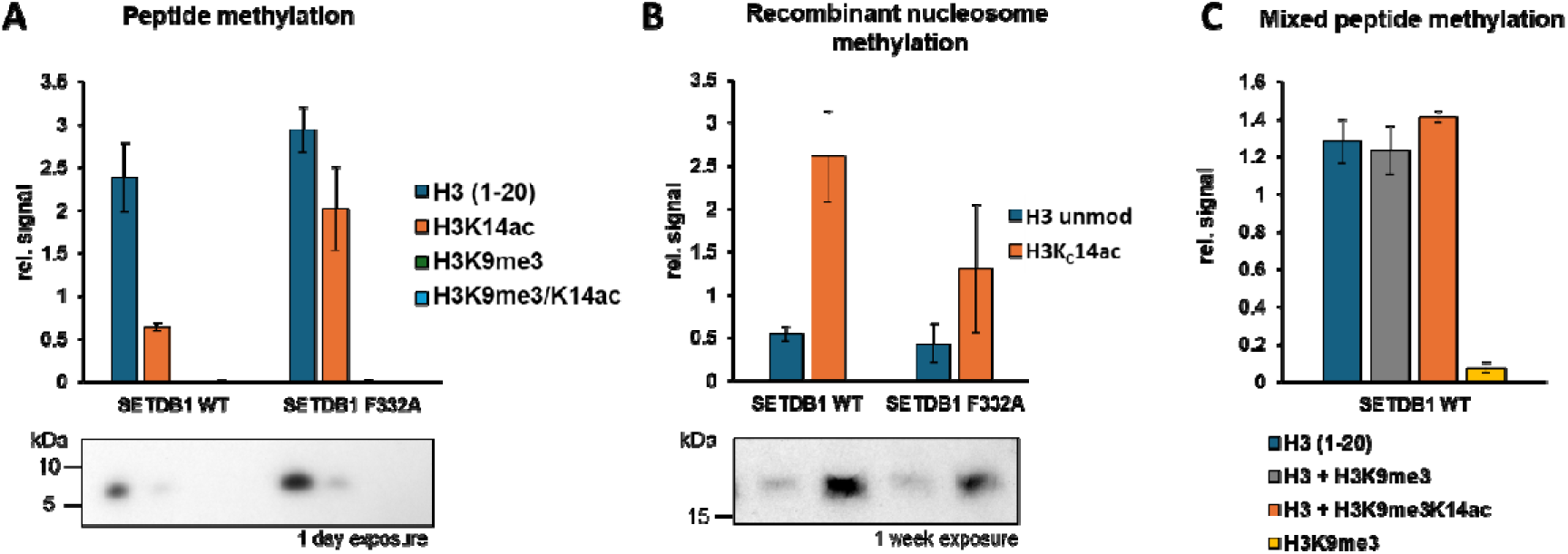
Peptide and recombinant nucleosome methylation by SETDB1. H3 (1-18) peptides (A), recombinant nucleosomes (B) mixed peptides (C) containing different modifications were incubated with full-length WT SETDB1 or its 3TD mutant F332A in methylation buffer containing radioactively labelled AdoMet and the transfer of methyl group was detected by autoradiography. The data are represented as means ± SEM of at least four (panel A) or three (panel B and C) independent experiments. See also Supplementary Figure 1 and 2.

In cells, SETDB1 methylates H3 tails on nucleosomes where two H3 tails are presented on the same face of the nucleosome suggesting that the SET and 3TD domains might interact with different tails of one nucleosome at the same time. To investigate this hypothesis, we prepared recombinant mononucleosomes either with unmodified H3 or H3 containing an acetyllysine analog at position 14 (H3K_C_14ac) (Supplementary Figure 3) ^36^. As shown in Figure 2B, H3K_C_14ac stimulates the H3K9 methylation activity of SETDB1 WT on the nucleosomal substrate by about 5-fold. In case of 3TD mutant F332A, we observed a reduction of the stimulation compared to SETDB1 WT. Taken together, the peptide and recombinant nucleosome data indicate that binding of 3TD to peptide containing H3K14ac inhibits H3K9 methylation of the same peptide, whereas in the nucleosome it stimulates SETDB1 activity for methylation of K9 on the other histone tail.

Two models could explain the stimulation of SETDB1 activity on the H3K14ac modified nucleosomes: In an allosteric model, H3K14ac peptide binding to the 3TD might trigger a conformational change of SETDB1 that leads to higher activity of the SET domain. Alternatively, in the recruitment model, 3TD binding to one H3 tail may help to position the SET domain close to the second tail leading to elevated methylation activity. In order to investigate if the binding of 3TD to K14ac can stimulate the catalytic activity of SETDB1 by inducing a conformational change, we performed mixed peptide methylation experiments, where one peptide could act as a substrate for 3TD binding and methylation of the other peptide by the SET domain is measured. As seen in figure 2C, addition of peptide containing H3K14ac-K9me3 which provides the 3TD binding motif but cannot be methylated itself did not stimulate the catalytic activity of SETDB1 (Supplementary Figure 2). These data argue in favor of the recruitment model suggesting that one H3K14ac tail acts as an anchor for 3TD binding and this directs the SET domain to methylation of the other histone tail. This process resembles the read/write mechanism of H3K9me3 propagation based on the Chromodomain and SET domain of SUV39H1 ^37^.

### SETDB1 is recruited to regions containing H3K14ac

Our enzyme kinetic data suggest that H3K14ac plays an important role in the recruitment of SETDB1. To investigate this hypothesis in cells, we generated a CRISPR-Cas9 mediated SETDB1 knock-out in HCT116 cells (Figure 3A, Supplementary Figure 4A). Afterward, the KO cells were rescued with SETDB1 WT, or SETDB1 containing either the F332A mutation in the 3TD or the H1224K mutation in the SET domain, which renders the enzyme catalytically inactive ^21^. All SETDB1 constructs were ectopically expressed under a constitutive promoter (Supplementary Figure 4D). We then performed Chromatin Immunoprecipitation (ChIP) of H3K9me3 followed by qPCR to analyze H3K9me3 levels in known SETDB1 target regions ^38,39^. As shown in Figure 3A, we observed a reduction of the original H3K9me3 levels (blue bars) in the SETDB1 KO cells (orange bars). H3K9me3 was effectively restored upon rescue with SETDB1 WT (green bars), however, it was not restored by the catalytically inactive mutant H1224K (pink bars). Interestingly, ChIP-qPCR results on SETDB1 targets showed differential recovery patterns by the F332A mutant depending on the target, where H3K9me3 at ERV3-I could not be effectively recovered, whereas it was partially recovered at ZNF221 and CDKL3 compared to parental HCT116 (blue bars) or SETDB1 WT rescue (green bars) (Figure 3B). This result suggests that an intact 3TD is required for efficient H3K9me3 methylation at most regions, which correlates with our in vitro experiments with nucleosomal substrates.

**Figure 3:**
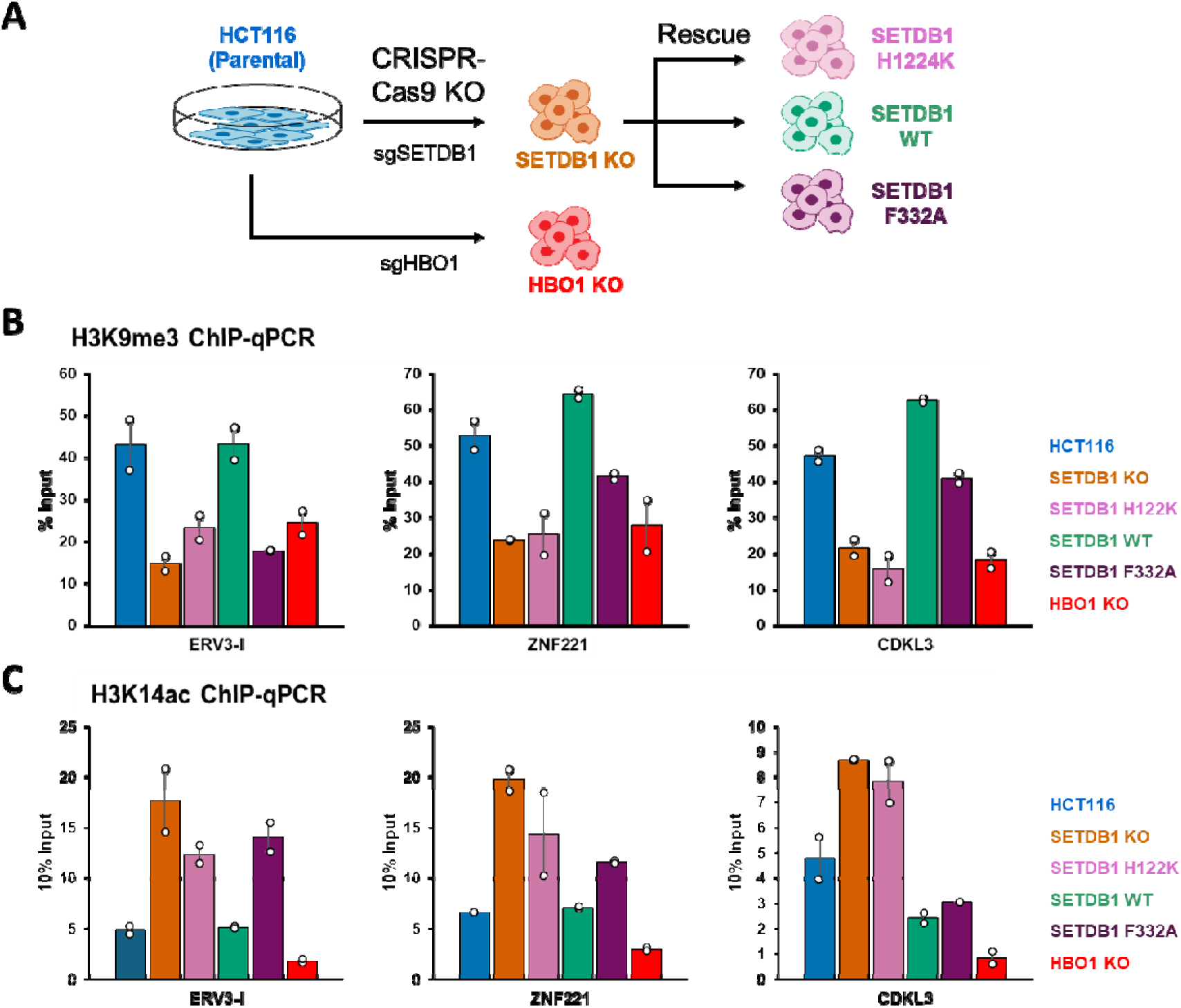
SETDB1 is recruited to H3K14ac containing target loci. **A)** Scheme of the cell HCT116 derived lines generated in this study by SETDB1 KO followed by reconstitution with SETDB1 WT or mutants, and by HBO1 KO. **B, C)** H3K9me3 (B) and H3K14ac (C) ChIP-qPCR at known SETDB1 target regions ^38,39^ showing changes in the histone modification as indicated upon SETDB1 KO and reconstitution with WT SETDB1 of its catalytically inactive mutant H1224K, or the 3TD mutant F332A. Moreover, HBO1 KO cells were investigated. ChIP was performed on mononucleosomes isolated from two individual biological replicates from each cell lines (represented as dots). Averages of the values are represented as bars.

We next performed H3K14ac ChIP-qPCR in the same regions and observed a robust H3K14ac signal in all regions (Figure 3C), in line with our hypothesis that H3K14ac plays an important role in SETDB1 targeting. Since SETDB1 is known to recruit the NuRD HDAC complex ^21^, we also investigated the changes in H3K14ac in SETDB1 KO cells and upon SETDB1 rescue. As shown in Figure 3C, H3K14ac levels in these regions increased drastically upon SETDB1 KO and they were brought to original levels after reconstitution with WT SETDB1, which illustrated the co-targeting of HDAC activity together with SETDB1. Interestingly, the recovery of H3K14ac after reconstitution with the H1224K mutant was incomplete suggesting that SETDB1 (and the associated NuRD complex) is only strongly recruited to the targets containing H3K14ac when H3K9 can be methylated by SETDB1, which agrees with the strong stimulation of 3TD binding to H3K9me1/2/3-K14ac when compared to H3K14ac alone ^23^. The recovery of H3K14ac was variable among the loci after reconstitution with SETDB1 F332A again indicating that the role of the 3TD in SETDB1 recruitment is locus dependent.

### Genome-wide analyses of SETDB1 activity

To explore the SETDB1 dependent dynamics of H3K9me3 genome-wide, we performed H3K9me3 ChIP coupled with Next Generation Sequencing (ChIP-Seq). Raw reads were trimmed and mapped to the latest human genome build T2T-CHM13v2.0 ^40^. This genome build covers the complete genome annotated from telomere to telomere which is especially important, because SETDB1 is known to target RE. ChIP-seq experiments were conducted in two independent biological replicates. The binding profiles observed in both experiments showed excellent correlations (Supplementary Figure 5A) and were merged. Calling statistically highly significant peaks was conducted on both replicates and manually checked for accuracy and reliability (Supplementary Figure 5B).

Comparison of the H3K9me3 peaks in parental HCT116 and SETDB1 KO cells revealed that the majority of the genome-wide H3K9me3 peaks (72% of all 153,144 peaks) were established by SETDB1 (Figure 4A, 4B and Supplementary Figure 6A). Knockout of SETDB1 led to loss of the H3K9me3 signal in the SETDB1 dependent cluster but an increase in H3K9me3 signal in SETDB1 independent cluster suggesting some compensatory mechanisms. Of note, highly repetitive heterochromatin, which is a bona fide SUV39H1/H2 target ^41^ is underrepresented in these data due to lower recovery of this chromatin from cells and low mappability. The majority of the peaks specifically lost upon SETDB1 KO were recovered after reconstitution with SETDB1 WT but not with the inactive H1224K SET domain mutant indicating that the methylation is indeed introduced by the catalytic activity of SETDB1 (Supplementary Figure 6A).

**Figure 4:**
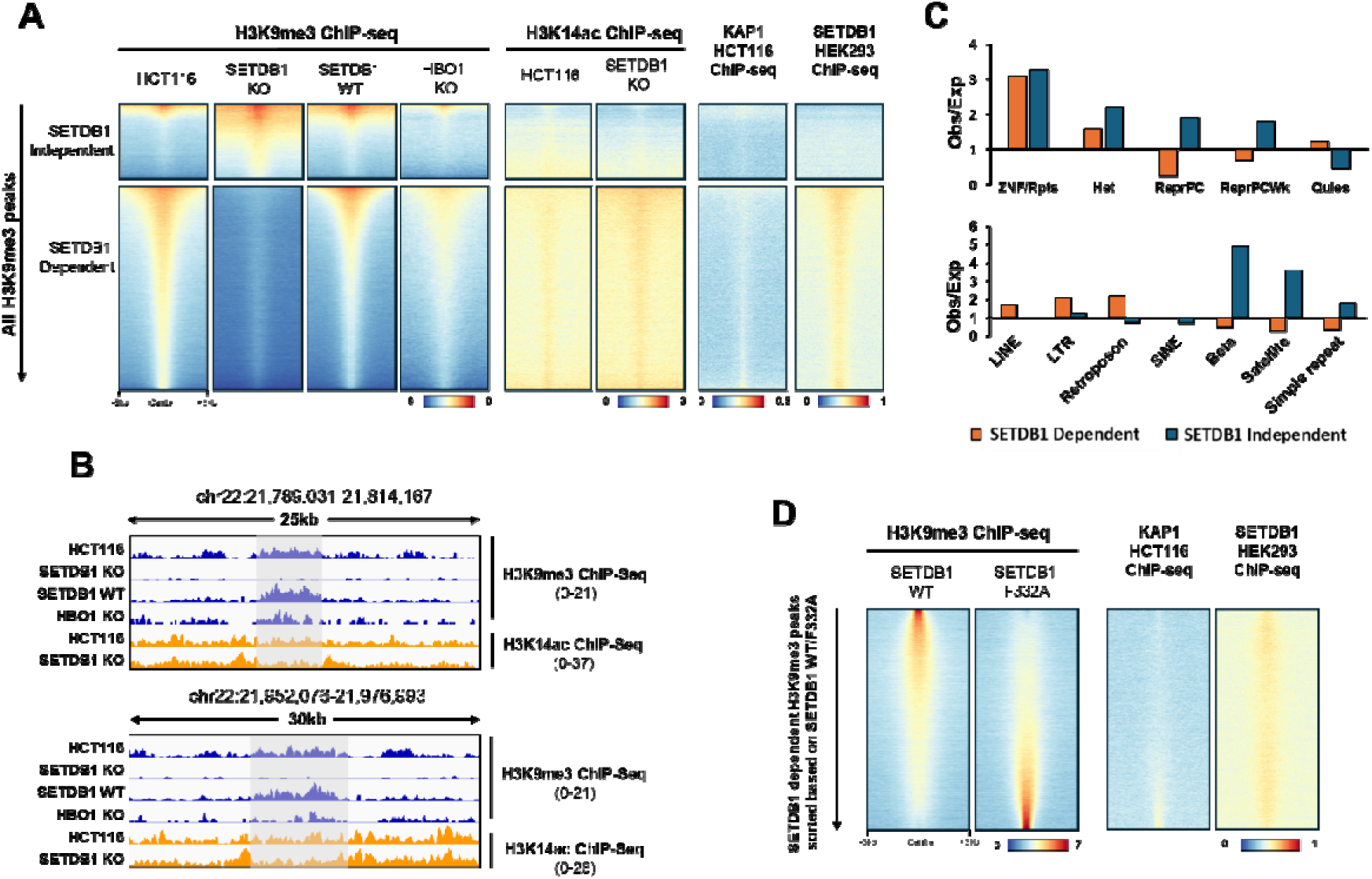
SETDB1 is dependent on H3K14ac for efficient methylation of H3K9. **A)** Heatmaps were generated using all H3K9me3 peaks from the different cell lines for H3K9me3 and H3K14ac ChIP-seq as indicated and were plotted as regions with 3 kb flanking either side. K-means clustering was based on the differential H3K9me3 signals, either as SETDB1 independent or SETDB1 dependent. Sorting was by decreasing H3K9me3 intensity. Datasets form the literature of KAP1 in HCT116 cells (GSM1866695) ^44^ and SETDB1 in HEK293 cells (GSM5331059) ^45^. **B)** IGV browser views of ChIP-seq profiles at exemplary SETDB1 dependent regions. **C)** Enrichment and depletion of SETDB1 dependent and independent H3K9me3 peaks in different chromatin regions. In upper panel, occurrence of peaks in chromatin segmentation (ChromHMM) of HCT116 cells was determined and compared with randomized peaks to calculate observed/expected ratios. In the lower panel, occurrence of peaks in RE annotated by RepeatMasker was analyzed in the same way. ZNF/Rpts: Zinc-finger genes and RE, Het: Heterochromatin, ReprPC: repressed PolyComb, ReprPCWk: Weak repressed PolyComb, Quies: Quiescent. **D)** Heatmap of H3K9me3 ChIP-seq in SETDB1 KO cells after rescue with WT or F332A SETDB1. SETDB1 dependent H3K9me3 regions were sorted by decreasing intensity of SETDB1 WT/F332A, which was then applied for sorting KAP1 and SETDB1 tracks.

We analyzed the distribution of SETDB1 dependent and independent peaks in different ChromHMM regions of HCT116 cells using ENCODE data (ENCODE ENCFF513PJK) and in regions annotated as RE by RepeatMasker (http://www.repeatmasker.org) (Figure 4C). We observed a depletion of SETDB1 dependent peaks in the regions annotated as PolyComb repressed, which is in agreement with the observation that H3K9me3 and H3K27me3 represent alternative mechanisms of gene silencing and heterochromatin formation ^42^. Overlap of H3K9me3 peaks with repeat masker annotated RE revealed enrichment of SETDB1 dependent peaks in LINE, LTR and retrotransposons, in line with the known role of SETDB1 in silencing these types of RE ^6^.

We also analyzed published H3K9me3 ChIP-seq data from SETDB1 WT and KO HeLa cells along with ChIP-seq of SETDB1 complex partners like HUSH (TASOR, MPP8 and periphilin) ^15^ and ATF7IP (Supplementary Figure 6B) ^38^. We mapped the raw data to T2T-CHM13v2.0 human genome and clustered it similar to Figure 4A. Knockout of SETDB1 led to loss of the H3K9me3 signal in the SETDB1 dependent cluster and an increase in H3K9me3 signal in SETDB1 independent cluster in this data set as well, showing that our clustering into SETDB1 dependent and independent regions is applicable in HeLa cells as well. ATF7IP KO showed an almost complete loss of H3K9me3 in the SETDB1 dependent regions similar to SETDB1 KO. Interestingly, KO of HUSH complex subunits had only a small effect on global H3K9me3. These findings underscore the particular importance of ATF7IP in stabilizing and recruiting SETDB1 ^38,43^.

### Effect of H3K14ac on SETDB1 recruitment

We next aimed to explore the effect of H3K9me1/2/3-K14ac binding by 3TD on the recruitment of SETDB1, but genome-wide ChIP-seq analysis of the different H3K9me1/2/3-K14ac dual marks is technically not possible. However, quantitative mass spectrometric histone modification data from HCT116 cells revealed that H3K9me1/2/3 is an extremely abundant modification and 88% of all histone tails containing H3K14ac also carry H3K9me1/2/3 ^25^. Therefore, H3K14ac alone can be studied as a proxy for H3K9me1/2/3-K14ac and we performed H3K14ac ChIP-seq in parental HCT116 and SETDB1 KO cells. Using the same H3K9me3 peak-based clustering for H3K14ac tracks, we observed a co-localization of H3K14ac mainly on SETDB1 dependent K9me3 peaks illustrating a genome-wide role of H3K14ac in SETDB1 targeting (Figure 4A). Moreover, we observed an increase in H3K14ac signals upon SETDB1 KO which can be associated with the lack of HDAC co-recruitment to these sites after SETDB1 KO. This finding further implies that these regions were even more enriched with H3K14ac before SETDB1 mediated introduction of H3K9me3 and HDAC induced H3K14ac deacetylation. We also analyzed public ChIP-seq data for KAP1 from HCT116 cells ^44^ and SETDB1 from HEK293 cells ^45^ again using the same H3K9me3-based clustering, and observed a very good colocalization of SETDB1 specifically on the SETDB1-dependent H3K9me3 peaks along with its known interactor KAP1 ^21^. In untreated cells, the SETDB1 independent H3K9me3 peaks showed very low levels of SETDB1 and KAP1 in addition to the reduced levels of H3K14ac.

To investigate the role of the 3TD on SETDB1 activity in more detail, the SETDB1 dependent peaks from SETDB1 WT and F332A reconstituted cells were used and sorted by the relative H3K9me3 WT/F332A RPKM ratio (Figure 4D). Arranged in the same order, KAP1 and SETDB1 ChIP-seq data were displayed. While SETDB1 was present at all regions, the data clearly illustrate that regions in which H3K9me3 reconstitution worked best with the F332A mutant contain more KAP1 suggesting that 3TD independent regions are more dependent on KAP1 for SETDB1 targeting.

### SETDB1 activity at ZNF genes

SETDB1 is known to be recruited to 3’ ends of ZNF genes through KRAB zinc finger containing binding protein ZNF274 and KAP1 ^46^. Hence, we looked at the ZNF genes which were sorted and clustered based on H3K9me3 levels in parental HCT116 and SETDB1 KO cells. As shown in Supplementary Figure 7, in the SETDB1 targeted ZNF clusters, H3K9me3 is established in the promoters of ZNF genes and the bodies peaking at the 3’ ends, and this H3K9me3 signal is lost upon SETDB1 KO. Similarly, SETDB1 is enriched at the promoter and the 3’ end of ZNF genes. SETDB1 enrichment at the promoters of ZNF genes along with KAP1 was co-occurring with H3K14ac. At these sites, H3K14ac increased on SETDB1 KO, again suggesting that SETDB1 is recruited to the region containing H3K14ac where it then brings in HDAC activity. K-means clustering of H3K9me3 levels showed a second cluster of ZNF genes which are depleted for SETDB1 and show low H3K9me3 levels despite having high levels of H3K14ac and KAP1 indicating that recruitment of SETDB1 is precluded by a so far unknown signal.

### SETDB1 dependent changes in DNA methylation

SETDB1 has been previously shown to interact and colocalize with the DNA methyltransferase DNMT3A ^22^. Furthermore, loss of SETDB1 has been shown to result in reduction of DNMT protein levels ^47^. Therefore, we determined the DNA methylation changes upon SETDB1 KO in two independent biological replicates using the Infinium EPIC array. Differentially methylated loci were identified using the Qlucore Omics Explorer (ver. 3.6) by applying T-test statistics (q<0.05) and filtered for relevant methylation changes (variance>0.4). By this approach, we identified 3979 hypomethylated and 6612 hypermethylated CpG loci in SETDB1 KO cells as compared to control samples (Supplementary Figure 8A). Comparison with ChIP-seq data revealed that SETDB1 dependent loss of H3K9me3 is correlated with the reduction of DNA methylation. The regions showing reduced DNA methylation after SETDB1 KO also showed an enrichment of H3K14ac and SETDB1, suggesting that the loss of DNA methylation is dependent on the lack of SETDB1 recruitment to the H3K14ac rich regions in the SETDB1 KO cells (Supplementary Figure 8B).

### H3K14 acetylation increases SETDB1 mediated H3K9 trimethylation

Our genomic data presented so far indicate that there is a correlation of SETDB1 recruitment to the H3K14ac rich regions, followed by the establishment of H3K9me3 by SETDB1 and removal of acetylation by the co-recruited NuRD HDAC complex. Next, we wanted to directly document the mechanistic role of H3K14 acetylation in regulating SETDB1 mediated H3K9 trimethylation. Previous work has shown that HBO1 is the main enzyme to introduce H3K14ac ^48^. Therefore, we generated a CRISPR-Cas9 mediated HBO1 KO in HCT116 cells (Figure 3A) which was validated by Sanger sequencing (Supplementary Figure 4B and C). We then performed H3K9me3 and H3K14ac ChIP-qPCR at SETDB1 target regions and observed a significant loss of H3K14ac levels compared to parental HCT116 cells (Figure 3C, red *vs*. blue bars). In line with our hypothesis, we also detected a significant reduction of H3K9me3 levels in the SETDB1 target regions lacking H3K14ac in HBO1 KO cells (Figure 3B, red *vs*. blue bars). This finding confirms that the H3K9me3 and H3K14ac are not antagonistic marks but rather the presence of H3K14ac is required for SETDB1 mediated H3K9me3, which is also consistent with our in vitro biochemical data. In agreement with our findings, previous acetylome mass spectrometry study also showed that loss of H3K14ac after HBO KO was accompanied by a loss of H3K9me3 in the same peptide ^26^.

We next wanted to investigate the relationship of H3K14ac and H3K9me3 genome-wide and we performed H3K9me3 ChIP-seq in HBO1 KO cells. Applying the same clustering to SETDB1 dependent and independent H3K9me3 regions as described before, we observed a very strong reduction of H3K9me3 in the SETDB1 dependent regions in HBO1 KO cells (Figure 4A), in agreement with the H3K9me3 ChIP-qPCR results. The loss of H3K9me3 upon HBO1 KO is also shown in the IGV browser view in Figure 4B. This result further supports our previous conclusion that the presence of H3K14ac is required for SETDB1 mediated methylation of target H3K9. As 3TD is the only part of SETDB1 known to bind H3K14ac, this also indicates the broad contribution of 3TD to SETDB1 recruitment. Thereby, our data demonstrate that H3K9me3 and H3K14ac are not antagonistic marks but rather the presence of H3K14ac is a strong supportive signal for SETDB1 dependent establishment of H3K9me3. Specifically, H3K14ac must be seen as particular kind of acetylation, which does not have its key role in gene activation, in line with the finding that the HBO1 genomic binding showed no correlation with levels of transcription ^48^.

### 3TD is important for repression of L1M RE

An independent study showed the enrichment of SETDB1 dependent H3K9me3 over the L1 RE and treatment with TSA resulted in specific increase in H3K14ac on H3K9me3 containing peptides in PC9 cell line ^49^. Hence, we analyzed the genome-wide distribution of H3K9me3 peaks on RE for all cell lines prepared here, showing an enrichment of SETDB1 dependent H3K9me3 peaks on the LINE, LTR, and retroelements as previously reported targets for SETDB1 ^6^ (Figure 5A). The loss of H3K9me3 at LTR, LINEs, and retrotransposons upon SETDB1 KO was efficiently reverted by recovery with WT SETDB1. However, at LINE and more specifically L1 RE, efficient recovery was dependent on an intact 3TD domain (Figure 5A). This finding was confirmed by an analysis of the enrichment of H3K9me3 peaks at individual RE sub-families in SETDB1 KO cells after reconstitution with F332A or WT SETDB1 (Figure 5B). This analysis revealed that the F332A mutant showed reduced H3K9me3 recovery at many instances of LINE RE, in particular L1 which is among the most active LINE retroelements ^50^. The detailed H3K9me3 profiles over L1M elements containing SETDB1 dependent H3K9me3 peaks confirmed this finding, by showing that the F332A mutant failed to recover H3K9me3 at L1M open reading frames (Figure 5C). In agreement with our previous findings, a slight increase in H3K14ac is observed upon SETDB1 deletion in the same regions.

**Figure 5:**
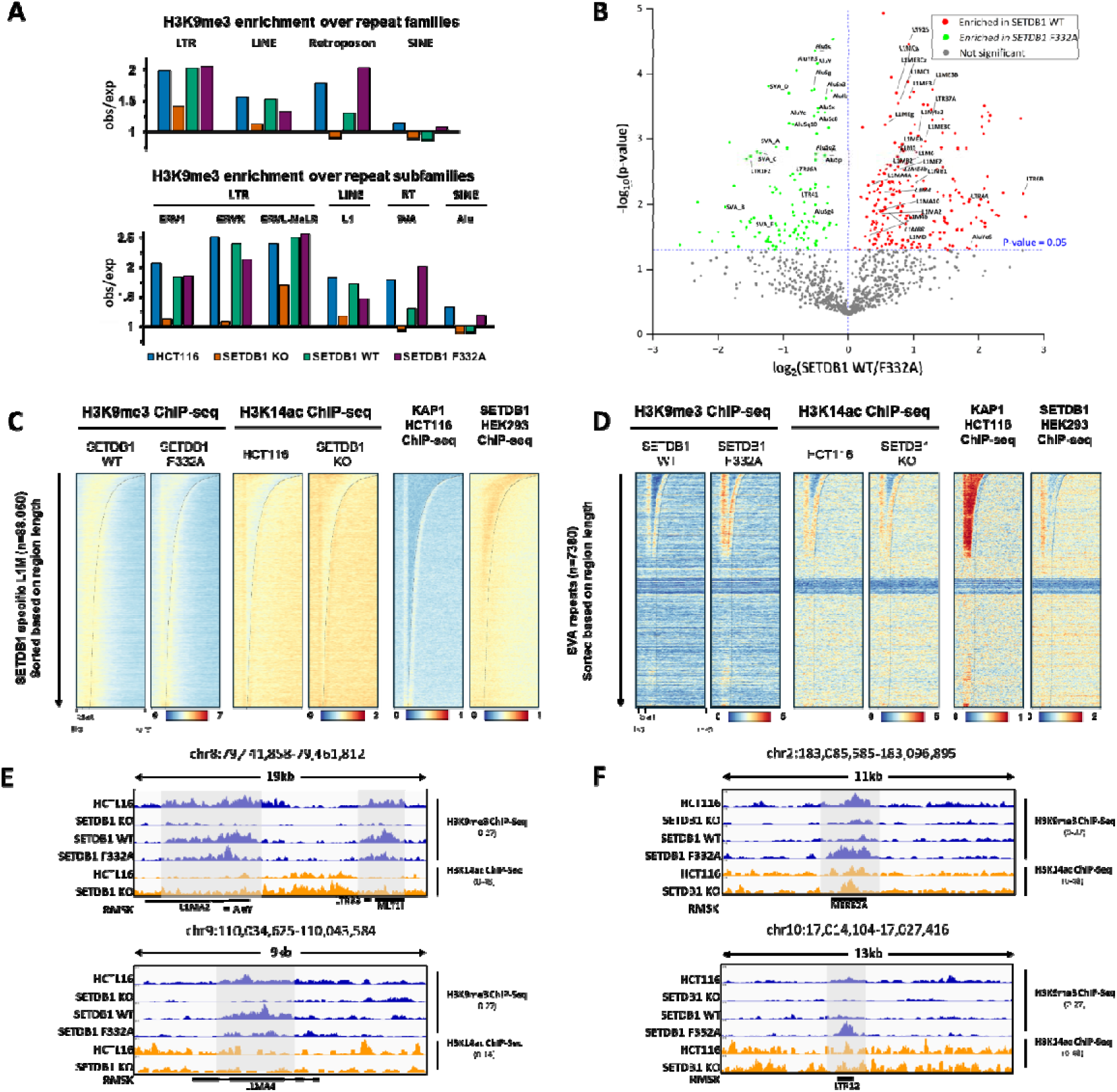
3TD of SETDB1 is important for the recruitment and efficient methylation in the L1M RE. **A)** Enrichment and depletion of H3K9me3 peaks of different cell lines in RE. RE families and subfamilies were annotated by RepeatMasker. **B)** Enrichment of H3K9me3 peaks in SETDB1 WT and F332A mutant cells for individual RE classes. The Log2-fold change of H3K9me3 peak enrichment in SETDB1 WT over F332A mutant is plotted against the corresponding p-value. Red dots represent individual RE with stronger methylation in SETDB1 WT. Some of the L1M elements which are only methylated in SETDB1 WT are annotated. Green dots represent RE with stronger methylation in the F332A mutant cell line showing enrichment of SVA and Alu elements. **C, D)** Heatmap H3K9me3 and K14ac signals on SETDB1 specific L1M RE (C) or SVA (SINE-VNTR-Alus) retrotransposons (D) sorted based on decreasing RE region length. **E, F)** Example browser views of H3K9me3 and H3K14ac tracks on representative L1M (E) and SVA (F) loci.

L1M RE are the evolutionarily oldest class of L1 elements ^17^. The young L1 elements like L1Ps were shown to be repressed by HUSH complex mediated recruitment of SETDB1 ^51^ which is driven by binding to long intron-less RNA ^17^. However, the HUSH complex is not enriched on the L1Ms, which do not show strong transcription. Similarly, KAP1 only shows slight enrichment over the 3’ and 5’ end of L1M whereas SETDB1 shows strong enrichment over the L1M body. Taken togehter, these data suggest that L1M RE are repressed by SETDB1 but recruitment is neither dependent on the HUSH complex nor KAP1. Instead, the 3TD domain of SETDB1 is involved in the recruitment to these loci in an H3K14ac dependent manner.

Differential H3K9me3 peaks of SETDB1 WT and F332A at the individual repeat classes showed that at some RE, for example SVA, H3K9me3 levels were even higher after KO recovery with 3TD than with WT SETDB1 (Figure 5A and B). The detailed H3K9me3 profiles over these SVAs containing SETDB1 dependent H3K9me3 peaks confirmed this finding, by showing that the rescue with F332A mutant was even more efficient than rescue with SETDB1 WT (Figure 5D). This could be explained by a high enrichment of KAP1 at these RE indicating that recruitment of SETDB1 to these RE is mediated by KAP1 and not dependent on 3TD. The increase in H3K9me3 reconstitution by the F332A mutant may be explained by increased activity of the mutant as observed in our in vitro biochemical data.

## Conclusions

In various cell lines about 25% and up to 50% of the H3 proteins were shown to contain the H3K9me1/2/3-K14ac modification ^25^ indicating that it is a very common double modification. This dual modification is bound by the SETDB1 3TD ^23^, but the relevance of this was unknown. Collectively our biochemical and genomics data indicate that SETDB1 is globally recruited to target regions through H3K14ac binding by 3TD where it leads to generation of H3K9me3 and H3K14ac deacetylation by the co-recruited NuRD HDAC complex (Figure 6). Combined with previous findings this illustrates that various mechanisms are involved in SETDB1 targeting, viz. KAP1 and HUSH complex recruitment and 3TD binding of H3K9me1/2/3-K14ac. The strong global correlation of H3K14ac with SETD1 binding and activity furthermore indicates, that 3TD binding cooperates at many genomic regions with the other targeting processes. However, our findings on RE illustrate that specific recruitment mechanisms are of relevance at some targets. At one group of RE, like SVAs, SETDB1 is mainly recruited via the KAP1 pathway and apparently independent of 3TD. On the contrary, at evolutionary old L1M elements, the 3TD domain of SETDB1 has a pivotal role in the recruitment of SETDB1.

**Figure 6:**
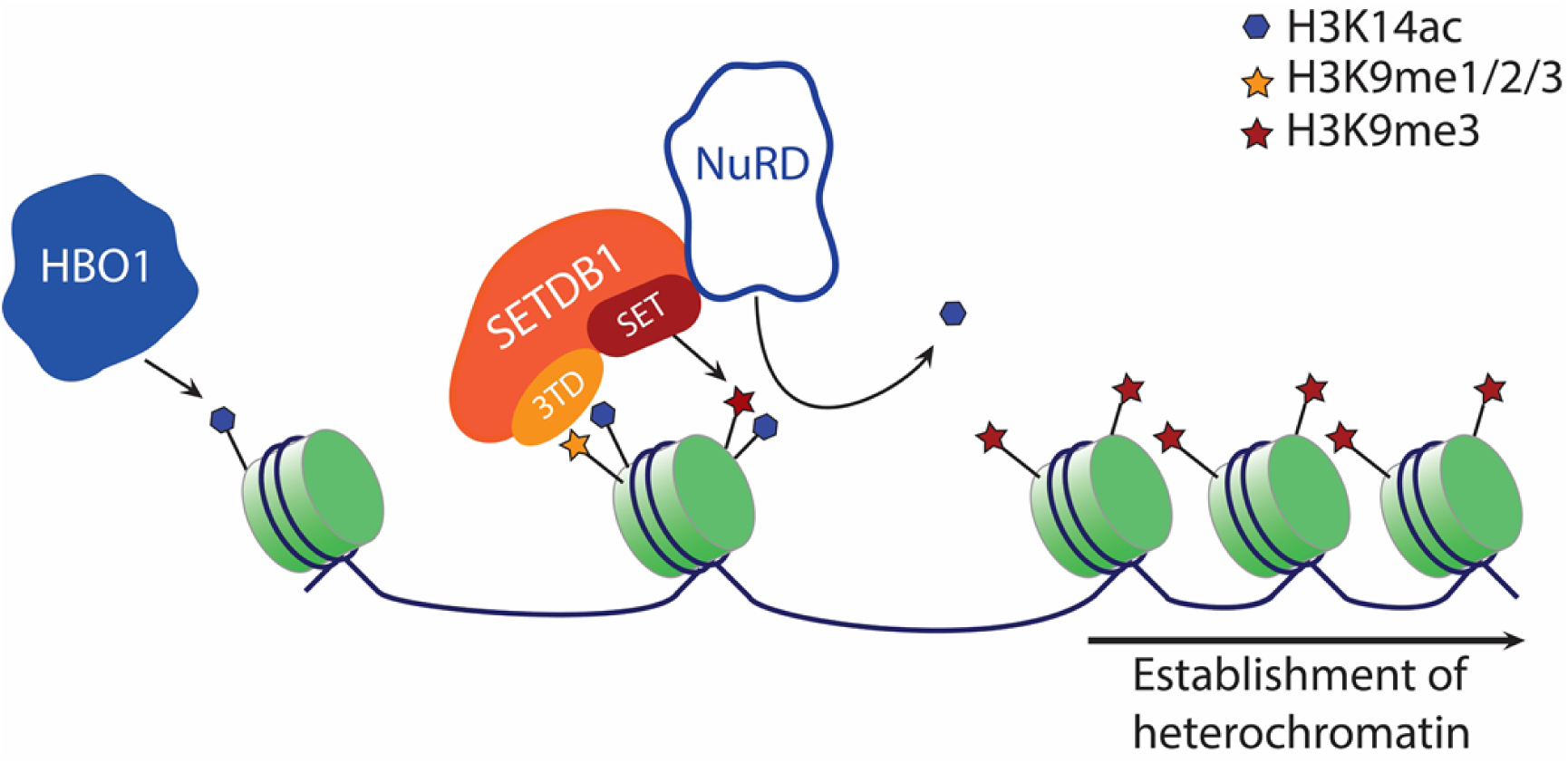
H3K14ac mediated efficient methylation of H3K9 by SETDB1. Schematic representation of the SETDB1 dependent trimethylation of H3K9 where 3TD is bound to H3K14ac, which is installed by HBO1, in one histone tail positioning SET domain of SETDB1 methylating the other H3K9 of the same nucleosome followed by the removal of H3K14ac by co-recruited NuRD complex resulting in the formation of heterochromatin.

### Materials and methods Cell culture

HCT116 cells were cultured in McCoy’s 5A (Sigma-Aldrich) media and Platinum-E (PlatE) retroviral packaging cell lines were cultured in DMEM high glucose media (Sigma-Aldrich) and both were supplemented with 10% Fetal Bovine Serum (FBS) (Sigma-Aldrich), 20 mM L-glutamate (Sigma-Aldrich) and 100 U/ml penicillin/streptomycin (Sigma-Aldrich). The cells were grown at 37°C and 5% CO_2_ and split every 2–3 days to keep confluency at 70-80%. For sub-culturing, the cells were washed with Dulbecco’s Phosphate Buffered Saline without MgCl_2_ and CaCl_2_ (Sigma-Aldrich), immersed with Trypsin-EDTA solution (Sigma Aldrich) and incubated at 37°C for 5 min. Trypsinization was stopped by resuspension in respective media and cells were usually split in a 1:4–1:8 ratio.

Sf9 insect cells were cultured in Sf-900 II SFM medium (Gibco), supplemented with 10% FBS at 25°C under suspension culture conditions at 150 rpm. Cell counts were determined every other day, and cells were sub-cultured twice weekly to ensure maintenance in the exponential growth phase. For transfection and viral plaque assays, cells from the suspension culture were seeded into 6 well plates at a density of 1 × 10^6^ cells per well.

### SETDB1 purification

Full-length SETDB1 (UniprotKB Q15047) and its F332A mutant were overexpressed and purified using Bac-to-Bac baculovirus expression system (Invitrogen) basically as described ^52^. In short, N-terminal His_6_- and C-terminal P2A separated EYFP full-length SETDB1 constructs were cloned into the pFastBacHT vector. The vector was then transformed into *E. coli* DH10Bac cells for the generation of recombinant Bacmid. Sf9 cells were transfected with the Bacmids using FuGENE, then followed by baculovirus amplification in Sf9 cells. Virus titer was determined by viral plaque assay. For protein expression, Sf9 cells were infected with the baculovirus with a multiplicity of infection of 6 at a cell density of 1X10^6^ cells/ml. Infected cells overexpressing full-length SETDB1 WT or F332A mutant were harvested 48 h after infection by centrifugation. Sf9 cells were then resuspended, lysed by sonication in buffer 1 (20 mM HEPES pH 7.2, 20 mM Imidazol, 500 mM KCl, 10% Glycerol, 0.5 mM DTT) supplemented with protease inhibitor cocktail (AEBSF-HCL (Biosynth), pepstatin (Roth), aprotinin and E-64 (Applichem), leupeptin and bestatin (Alfa Aesar)) and centrifuged for 1 h at 20,000 rpm at 4°C. The supernatant was transferred onto a Ni-NTA agarose bead (Qiagen) purification column, which was pre-equilibrated with buffer 1 followed by washing the protein bound beads with buffer 2 (20 mM HEPES pH 7.2, 20 mM Imidazole, 250 mM KCl, 10% Glycerol, 0.5 mM DTT). Bound proteins were then eluted in elution buffer (20 mM HEPES pH 7.2, 250 mM Imidazole, 250 mM KCl, 10% Glycerol, 0.5 mM DTT). Eluted proteins were dialyzed in dialysis buffer (20 mM HEPES pH 7.2, 250 mM KCl, 10% Glycerol, 0.5 mM DTT) for 2h and aliquots of full-length SETDB1 and F332A mutants were flash frozen in liquid nitrogen and stored at -80°C. The concentration was determined using Nano-Drop (Thermo Scientific) based on OD^280nm^ and SDS-PAGE was used to verify the concentration and purity of the proteins.

### Acetyl lysine analog installation, histone purification, and nucleosome reconstitution

Nucleosome reconstitution was performed as described ^53,54^. Briefly, pET21a expression constructs of H3.1, H4, H2A, and H2B were transformed into BL21-Codon Plus *E.coli* grown until OD^600nm^ of 0.6 to 0.8 at 37°C. Proteins were overexpressed by the addition of 1 mM IPTG and cells were grown for 3 h at 20°C. Cells were then harvested by centrifugation at 5000 x g for 15 min and washed using STE buffer (10 mM Tris/HCl pH 8, 100 mM NaCl, 1 mM EDTA), centrifuged again at 5000 x g for 15 min and the cell pellets were stored at - 20°C.

For histone protein purification, bacterial cell pellets were resuspended in SAU buffer (36 mM sodium acetate pH 7.5, 1 mM EDTA, 10 mM lysine, 5 mM β-mercaptoethanol, 6 M urea, 200 mM NaCl) followed by sonication (15 s pulse, 14 rounds, 30% power, 4°C) (Epishear, Active Motif). Cell lysates was then centrifuged at 40,000 x g for 1 h to remove the cell debris and supernatant was further filtered through 0.45 µm syringe filter (Chromafil GF/PET 45, Macherey Nagel). For the next step, a HiTrap SP HP (5 ml, GE Healthcare) column was connected to an NGC FPLC (BioRad) and equilibrated with SAU buffer before the injection of the filtered cell lysate onto the column. The column was then washed with SAU buffer and proteins were eluted with NaCl gradient from 200 mM to 800 mM. Fractions were collected, analyzed by SDS-PAGE, and pooled according to purity and yield. Finally, samples were dialyzed against pure water with two changes overnight and dried in a vacuum centrifuge for storage at 4°C.

Acetyllysine analog was generated as described using nucleophilic exchange of cysteine residues ^36^. The H3K14C cysteine mutation was introduced by site-directed mutagenesis. H3K14C proteins were dissolved in reaction buffer (200 mM sodium acetate buffer pH 4, 50 mM N-vinyl-acetamide, 5 mM VA-044, 15 mM reduced glutathione, 6 M guanidium chloride). The mixture was incubated at 20°C for 2 h in a cuvette placed in a Jasco FP-8300 spectrofluorometer under UV at 365 nm (slit width 5 nm). The converted histones were purified in PD-10 desalting column (GE Healthcare) and dialyzed against water for 2 h. Histones were then aliquoted, dried in vacuum centrifuge and stored at 4°C. Successful conversion of the cysteine into acetyllysine-analog (H3K_C_14ac) was confirmed by MALDI mass spectrometry and western blot using an H3K14ac antibody (Millipore-Sigma, 07-353).

For histone octamer reconstitution, purified dried histone proteins were dissolved in unfolding buffer (20 mM Tris/HCl pH 7.5, 7 M guanidinium chloride, 5 mM DTT) and their concentrations were determined based on OD^280nm^. The proteins were mixed in a ratio of 1 (H2, H4) to 2 (H2A, H2B). To reconstitute octamers, they were dialyzed against refolding buffer (10 mM Tris/HCl pH 7.5, 1 mM EDTA, 2 M NaCl, 5 mM β-mercaptoethanol) overnight with one buffer change. Reconstituted octamers were then purified by size exclusion chromatography using Superdex 200 16/60 PG column equilibrated with refolding buffer. Eluted fractions were collected, the fractions with highest purity and yield were pooled, and afterwards concentrated using Amicon Ultra-1 centrifuge filters (30 kDa cutoff, Merck Millipore). The purified octamers were then validated by SDS-PAGE, aliquoted, flash frozen in liquid nitrogen, and stored at -80°C.

For nucleosome reconstitution, the Widom-601 sequence ^55^ cloned into a TOPO-TA vector along with a linker sequence providing 64 bp linker DNA on the 5’ side of the core nucleosome and 29 bp on the 3’ side ^53^ was amplified by PCR and purified. DNA and histone octamers were mixed in different ratios between equimolar and 1.5-fold excess of octamer. The samples were then dialyzed in a Slide-A-Lyzer microdialysis device (ThermoFisher) against high salt buffer (10 mM Tris/HCl pH 7.5, 2 M NaCl, 1 mM EDTA, 1 mM DTT), which was continuously replaced by low salt buffer (same as high salt, but with 250 mM NaCl) over 24 h. Afterward, the samples were dialyzed overnight against storage buffer (10 mM Tris/HCl pH 7.5, 1 mM EDTA, 1 mM DTT, 20% glycerol), aliquoted, flash-frozen in liquid nitrogen, and stored at −80 °C. The DNA assembly in the nucleosomes was validated by an electro-mobility gel shift assay of the bound DNA.

### Peptide array methylation

Peptide arrays were synthesized on a cellulose membrane by SPOT synthesis method ^56^ using a Multipep peptide synthesizer (CEM). Each peptide spot contains approximately 9 mmol of peptide (Autospot Reference Handbook, Intavis), the successful synthesis of each peptide was confirmed by bromophenol blue staining ^31,32^. In addition, peptide synthesis was further confirmed by including internal positive and negative control peptides in all experiments. For the methylation of peptide arrays, they were initially incubated in the methylation buffer (50 mM Tris/HCl pH 9, 5 mM MgCl_2_, 5 mM DTT) for 5 min. Later, the peptide arrays were incubated in methylation buffer containing either SETDB1 WT or F332A mutant and 0.76 µM radioactively labelled AdoMet (PerkinElmer) for 4 h at 25°C. The arrays were then washed with a washing buffer (100 mM NH_4_HCO_3_ and 1% SDS). Finally, arrays were incubated for 5 min in amplify NAMP100V solution (GE Healthcare) and then exposed to the Hyperfilm^TM^ high performance autoradiography film (GE Healthcare) in the dark at −80°C for 24 to 48 h. The autoradiography films were developed with an Optimax Typ TR machine after different exposure times. Quantification of scanned images was conducted with ImageJ.

### Peptide and recombinant nucleosome methylation

For peptide methylation, equal concentrations of SETDB1 WT and F332A mutant were incubated with 4.4 µM of H3 peptides (Supplementary Table 1) in methylation buffer (50 mM Tris/HCl pH 9, 5 mM MgCl_2_, 5 mM DTT) supplemented with 0.76 µM radioactively labelled AdoMet (PerkinElmer) for 4 h at 25°C. The reactions were stopped by the addition of SDS-PAGE loading buffer and heating for 5 min at 95°C. Samples were then separated by Tricine-SDS-PAGE followed by incubation of the gel in amplify NAMP100V (GE Healthcare) for 1 h on a shaker. Then the gels were dried for 2 h at 70°C under vacuum. Transfer of radioactively labelled methyl groups were detected by autoradiography as described above.

For recombinant nucleosome methylation, normalized amounts of unmodified and H3K_C_14ac modified nucleosomes were incubated with SETDB1 WT or F332A mutant in the methylation buffer supplemented with 0.76 µM radioactively labelled AdoMet for 4 h at 25°C. The reactions were stopped by the addition of SDS-PAGE loading buffer and heating for 5 min at 95°C. Samples were separated in 16% SDS-PAGE and processed as described above.

### Generation of knockout cell lines

Single guide RNA sequences were adopted from ^15^ for targeting SETDB1 and from ^48^ for targeting HBO1. Single stranded oligos were ordered from IDT and annealed to their complementary oligos to result in double stranded DNA with 5’ single stranded overhangs complementary to the BbsI restricted Cas9 vector. Double stranded oligonucleotides were then ligated into the pU6-(BbsI)CBh-Cas9-T2A-mCherry plasmid backbone (provided by Ralf Kuehn, Addgene catalog number 64324) in the presence of BbsI HF restriction enzyme (NEB) and T4 DNA ligase (NEB) using golden gate assembly. Assembled products were transformed into XL1 blue *E. coli* cells by electroporation and plasmids were isolated using QIAamp DNA midi kit (QIAGEN). The Cas9-sgRNA plasmids were validated by BbsI restriction digestion and by Sanger sequencing. Four Cas9-sgRNA plasmids were pooled (150ng/µl) and were used to transfect HCT116 cells at 70% confluency using FuGENE® HD Transfection Reagent (Promega). On day three after transfection, mCherry positive cells were single cell sorted at high stringency using Sony cell sorter SH800S into 96 well plates containing McCoy’s 5A supplemented with 20% FBS. Single cells clones were expanded and selected. In the case of SETDB1, knockout clones were screened by western blot using a SETDB1 antibody (Abcam, ab107225). For HBO1, clones were screened by western blot using an H3K14ac antibody (Millipore-Sigma, 07-353) for loss of the H3K14ac mark when compared to parental HCT116 cells. Knockout clones were validated at the genomic level by Sanger sequencing, for which genomic DNA was isolated using QIAamp DNA mini kit (QIAGEN), and PCR amplified with primers located ∼500-600 bp away from the sgRNA target site. PCR products were then cloned into TOPO-TA vector (Invitrogen) and individual clones were Sanger sequenced.

### Retroviral transduction

For the rescue experiments of SETDB1 KO cells, full-length human SETDB1 and respective catalytically inactive mutant H1224K ^21^ and 3TD mutant F332A with reduced H3K9me1/2/3-K14ac peptide binding ^23^ were cloned into the pMSCV-PGK-NEO-IRES-GFP vector ^57^ and 20 µg of plasmid were precipitated as described ^58^. Briefly, HCT116 SETDB1 KO cells were precipitated for 20 min in HBS buffer (140 mM NaCl, 25 mM HEPES, 0.75 mM Na_2_HPO_4_, pH 7.0) together with 125 mM CaCl_2_ and 10 μg pCMV-Gag-Pol (Cell Biolabs) helper plasmid. The mix was added to a 10 cm dish with Platinum-E cells growing at 75–85% confluence in supplemented DMEM. After 16 and 24 h, the media was replaced with fresh DMEM. Supernatant containing the virus was collected 40–50 h after transfection, filtered through a 0.45 μm filter and added to the target cells at 50–70% confluence. Stably integrated cell populations were enriched using antibiotic selection with 10 µg/ml of G418 for 10 days. The efficiency of stable integration of pMSCV vector was analyzed by FACS.

### Histone extraction

Native histones were extracted as described ^59^. Briefly, 1 x 106 cells were harvested and lysed using hypotonic lysis buffer (10 mM Tris/Cl pH 8.0, 1 mM KCl, 1.5 mM MgCl_2_ and 1 mM DTT) at 4°C under constant rotation. Intact nuclei were then pelleted by centrifuging at 10,000 x g for 10 min at 4°C. Supernatant was discarded and the pellet resuspended in 0.4 N H_2_SO_4_ and incubated for 2 h. Cell debris was pelleted. Afterward, the dissolved histones were transferred to fresh tube and then precipitated using 100% TCA. Histones were then pelleted at 16,000 x g, washed with ice cold acetone and air dried. Histones were dissolved in water and stored at -20°C until further use.

### Chromatin-Immuno precipitation

1 x 10^6^ cells were used for H3K9me3 ChIP and 1 x 10^7^ were used for H3K14ac ChIP. Both ChIP experiments were conducted similarly until otherwise stated and methods are described for H3K9me3 and were usually scaled 10X for the H3K14ac ChIP. Cells were lysed with 50 µl of lysis buffer (10 mM Tris/HCl pH 7.4, 2 mM MgCl_2_, 0.6% IGEPAL® CA-630, 0.5 mM PMSF, 1 mM DTT) including 100 μl/ml EDTA-free protease inhibitor cocktail (Roche, dissolved in 1 ml H_2_O) and incubated on ice. Samples were then digested with 135 U of MNase (NEB, M0247) per 1 x 10^6^ cells at 37°C for 10 min. Digestion was stopped by the addition of 6.6 μl of 100 μM EDTA. 6.6 μl of 1% Triton X-100/1% deoxycholate solution was added and the sample was kept on ice for 15 min. Samples were then diluted with 200 µl of Complete Immunoprecipitation buffer (20 mM Tris/HCl pH 8.0, 2 mM EDTA, 150 mM NaCl, 0.1% Triton X-100. 1 mM PMSF, containing 100 μl/ml EDTA-free Protease Inhibitor Cocktail (Roche). Samples were incubated for 1 h at 4°C under constant rotation. Samples were then vortexed and centrifuged for 10 min at 14,000 x g at 4°C. DNA concentration was determined by OD^260nm^ measured using Nano-Drop (Thermo Scientific).

25 µg of the isolated chromatin was used for the H3K9me3 and 95 µg was used for H3K14ac pull-down. For pre-clearing of chromatin, 20 µl of Protein A/G magnetic beads (60 µl in case of H3K14ac IP) were washed three times with 200 µl of Complete Immunoprecipitation buffer and 2.5 µg of Rabbit IgG (R&D Systems, #AB-105-C) was added and incubated with rotation for 2 h at 4°C. 10% of the chromatin was taken aside as input. H3K9me3 was precipitated using the direct pull-down method where chromatin was added to the antibody-bead complex prepared by washing 20 µl of Protein A/G magnetic beads followed by incubating with 2.5 µg H3K9me3 antibody (Diagenode, C15200146) for 3 h at 4°C under constant rotation. Afterwards, supernatant was removed and chromatin was transferred to the antibody-bead complex and then incubated overnight at 4°C under constant rotation. In case of H3K14ac, the indirect pull-down method was used, where chromatin was directly incubated with 10 µl of the H3K14ac antibody (Millipore-Sigma, 07-353) under the same conditions. Afterward, 60 µl of pre-washed Protein A/G magnetic beads were added to trap the antigen-antibody complex for 2 h at 4°C under constant rotation. Later, both samples were washed, initially two times with 400 µl of low salt buffer (20 mM Tris/HCl pH 8.0, 2 mM EDTA, 150 mM NaCl, 1% Triton X-100, 0.1% SDS), followed by two times high salt buffer (20 mM Tris/HCl pH 8.0, 2 mM EDTA, 500 mM NaCl, 1% Triton X-100, 0.1% SDS), each for 10 min at 4°C. For the H3K14ac pull-down, two additional washes with TEN buffer (20 mM Tris/HCl pH 8.0, 2 mM EDTA, 150 mM NaCl) were performed. After the washing steps, the supernatant was removed and 200 µl of ChIP elution buffer (1% SDS, 100 mM NaHCO3) as well as 2.4 U proteinase K (NEB) were added to the beads and also the Input chromatin was treated the same and incubated for 2 h at 65°C. The supernatant of the samples containing the released DNA fragments was separated from the beads and transferred into a fresh tube. DNA from the IP and Input was purified using ChIP DNA Purification Kit (Active Motif).

### Library preparation for ChIP-seq

Two biological replicates were prepared for each ChIP-seq experiment. Library preparation was carried out using NEBNext® Ultra™ II DNA Library Prep Kit following the manufacturer′s protocol. DNA quality and concentration was measured using LabChip® GXII Touch™ HT system (Perkin Elmer) and 15 to 27 µg of DNA were used. After end repair and adapter ligation, DNA was amplified using NEBNext® Multiplex Oligos for Illumina® (Index Primers Set 1 or 2) (NEB). The quality of the libraries was checked using LabChip® GXII Touch™ HT system (Perkin Elmer) sequenced on an Illumina NovaSeq 6000 with 150 bp paired-end reads for a minimum of 10 million reads.

### ChIP-seq data analysis

Sequencing data was received in FASTQ format and processed on a local instance of Galaxy server ^60^. The quality of the raw reads was assessed using *FastQC* and adaptors were trimmed using *Trim Galore!* with default settings. High-quality reads were mapped to the latest version of human genome build T2T-CHM13 v2.0 using *HISAT2* with --no-spliced-alignment set to true ^61^. The resulting bam files were used to calculate genomic coverage using *bamcoverage* in 25 bp bins and normalized by reads per kilobase per million mapped reads (RPKM). Reproducibility of the replicates was determined using *multiBigWigSummary* tool in 5 kb bins. Replicates were then merged using *MergeSamFiles* and converted again to BigWig files which were then used for plotting heatmaps and visualized in Integrative Genomics Viewer (IGV) ^62^. Raw ChIP-seq data of H3K9me3 ChIP-seq from HeLa cells (GSE63116), H3K9me3 ChIP-seq ATF7IP KO (GSM2308950) KAP1 (GSM1866695) and SETDB1 (GSM5331059) was downloaded from European Nucleotide Archive and was processed as mentioned above.

### Peak calling and heatmap

BAM files of biological replicates were pooled in BAMPE format with effective genome size of 3.03 x 10^9^. Peaks were called using *MACS2* with input as control using -broad regions. For heatmap and clustering, H3K9me3 peaks of all the samples were merged and the resulting BED file was used as regions to plot the intensities using the *computeMatrix* tool and perform *k*-means clustering. Heatmaps were generated using *plotHeatmap*, the sorting of the heatmap is indicated for each image.

### Chromatin segment and repeat analysis

The 18-state ChromHMM data for HCT116 cells were obtained from ENCODE (ENCFF513PJK). It was lifted to T2T-CHM13 v2.0 using UCSC lift genome annotations tool and repeatmasker file was downloaded from UCSC genome browser. RE were annotated using the family and sub-family assignment in the repeat masker file, and the ratio of observed to expected overlaps was calculated for the various RE families. Overlaps between the ChromHMM or repeat regions and the H3K9me3 peaks were calculated using the *AnnotateBed* function of *bedtools*. For control comparisons, peak files were randomized using *shuffleBed* maintaining the number and length of peaks.

### ChIP-qPCR

To validate the ChIP-seq experiments, ChIP-qPCR was performed for SETDB1 target regions taken from literature (Supplementary Table 2 and 3). qPCR was performed on two independent biological replicates. For that, 1:5 dilution series of the input DNA were prepared for normalization of the pulled-down DNA and evaluation of the PCR efficiency. For each region of interest, a master mix was prepared with 7.5 µl of 2X ORA™ See qPCR Probe Mix (highQu), 0.4 μl forward primer, 0.4 μl reverse primer and 5.7 μl ddH_2_O. Plate was setup with 1 µl of sample in triplicates for input dilution series, IP, and non-template control (NTC). PCR was run with the following program: 95°C for 3 min, 40 cycles of 95°C for 3 s, 57°C for 20 s, 72°C for 4 s and finally a 65–95°C ramp (0.5°C steps every 5 s). Data was analyzed in Bio-Rad CFX Maestro software. Quality of the qPCR run was assessed by checking single melting curve indicating the amplified product.

### DNA methylation analysis

Genomic DNA was extracted using QIAmp DNA Mini Kit (Qiagen). The EZ DNA methylation kit (Zymo Research, Irvine, CA, USA) was used to perform bisulfite conversion on a total of 1000 ng of genomic DNA following the manufacturer’s guidelines. DNA methylation in parental HCT116 and SETDB1 KO HCT116 was investigated using the Infinium MethylationEPIC BeadChip (EPIC, Illumina Inc., San Diego, CA, USA) following the manufacturer’s instructions. Raw intensity files (idat) were brought into the R programming environment (version 4.3.1) utilizing the minfi package ^63^. The data were normalized according to controls using the preprocess Illumina function without applying background correction. The levels of methylation at each CpG site were calculated as a beta value which ranged from 0 (no methylation) to 1 (complete methylation). Differentially methylated loci were determined (T-test q<0.05, variance>0.4) and subsequently presented in a heatmap using the Qlucore Omics Explorer ver. 3.6 (Qlucore, Lund, Sweden). Results were normalized (mean=0 and variance=1) and presented as z-scores (blue: low scores, yellow: high scores).

### Statistics and reproducibility

The number of independent experimental repeats is indicated for each experiment. Standard deviations and SEM were determined with MS Excel. P-values were determined using two-sided T-test assuming unequal variance with MS Excel.

## Data availability

All biochemical data generated or analyzed during this study are included in the published article and its supplementary files. The H3K9me3 and H3K14ac ChIP-seq data are available at GEO under the accession number GSE261744. DNA methylome data of HCT116 and SETDB1 KO HCT116 are available at GEO under the accession number GSE264388.

## Supporting information

This article contains supporting information.

## Author contributions

AJ devised and supervised the study. TC conducted the biochemical experiments with support from AB, SW and PB. TC conducted the bioinformatic analysis with support from MC and PB. TC conducted the DNA methylation analysis with the support from AK and OA. PR provided research materials and advice. All authors were involved in data analysis and interpretation. TC and AJ prepared the figures and wrote the manuscript draft. The final manuscript was approved by all authors.

