## Supplemental Information for "SETDB1 activity is globally directed by H3K14 acetylation via its Triple Tudor Domain"

### Supplementary Information

#### Supplementary Figures

Supplementary Figure 1: Additional data regarding SETDB1 proteins and peptide SPOT array methylation.

Supplementary Figure 2: Mixed peptide methylation by SETDB1.

Supplementary Figure 3: Purification and reconstitution of recombinant nucleosomes.

Supplementary Figure 4: Validation of SETDB1 and HBO1 KO cell clones in HCT116 cells.

Supplementary Figure 5: Quality check NGS data analysis.

Supplementary Figure 6: Additional ChIP-seq analysis.

Supplementary Figure 7: H3K9me3, H3K14ac, KAP1 and SETDB1 ChIP-seq of different cell lines.

Supplementary Figure 8: SETDB1 dependent changes in DNA methylation.

#### Supplementary Tables

Supplementary Table 1: List of peptides used in this study.

Supplementary Table 2: List of sgRNA used for CRISPR/Cas9 mediated knockout in this study.

Supplementary Table 3: List of ChIP-qPCR primers and amplicons used in this study.

#### Supplementary References

**Supplementary Figures**

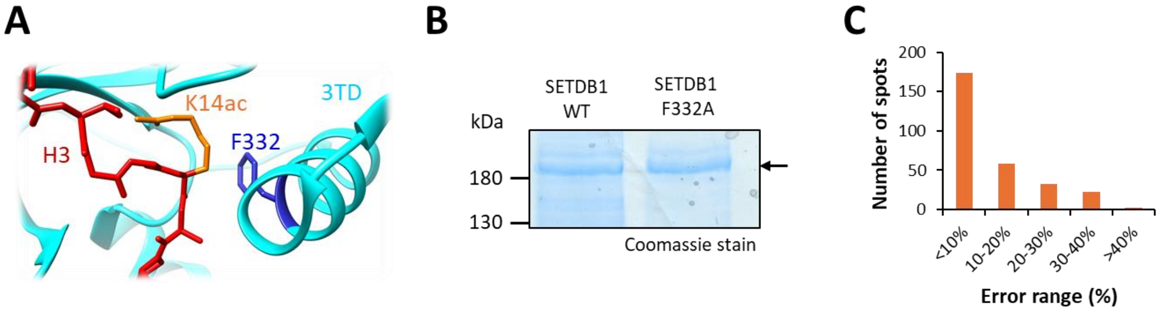


**Supplementary Figure 1: Additional data regarding SETDB1 proteins and peptide SPOT array methylation. A)** Structural snapshot showing the hydrophobic interaction of F332 and K14ac. 3TD is shown in cyan ribbon with F332 in blue. The H3 peptide is shown in red with the K14ac side chain in orange. **B)** Full-length purified protein of SETDB1 and 3TD mutant F332A were normalized and loaded in 16% SDS gel. The band corresponding to SETDB1 is indicated by an arrow. **C)** Error distribution of specificity array analysis shown in Figure 1 calculated by standard deviation of intensities of corresponding spots in both the replicate experiments.

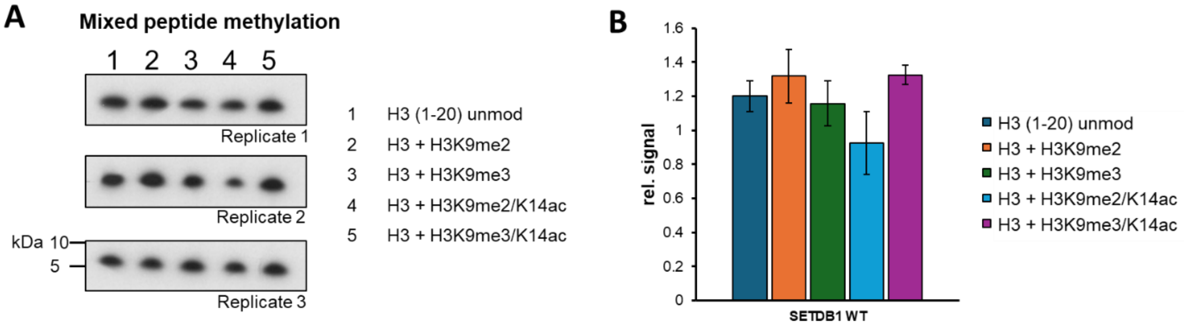


**Supplementary Figure 2: Mixed peptide methylation by SETDB1. A)** Autoradiography image of the mixed peptide methylation experiments shown in Figure 2C also including the corresponding H3K9me2 data. **B)** Averages of the results of the full panel of mixed peptide methylation experiments. Experiments were conducted in triplicate and the corresponding SD are shown.

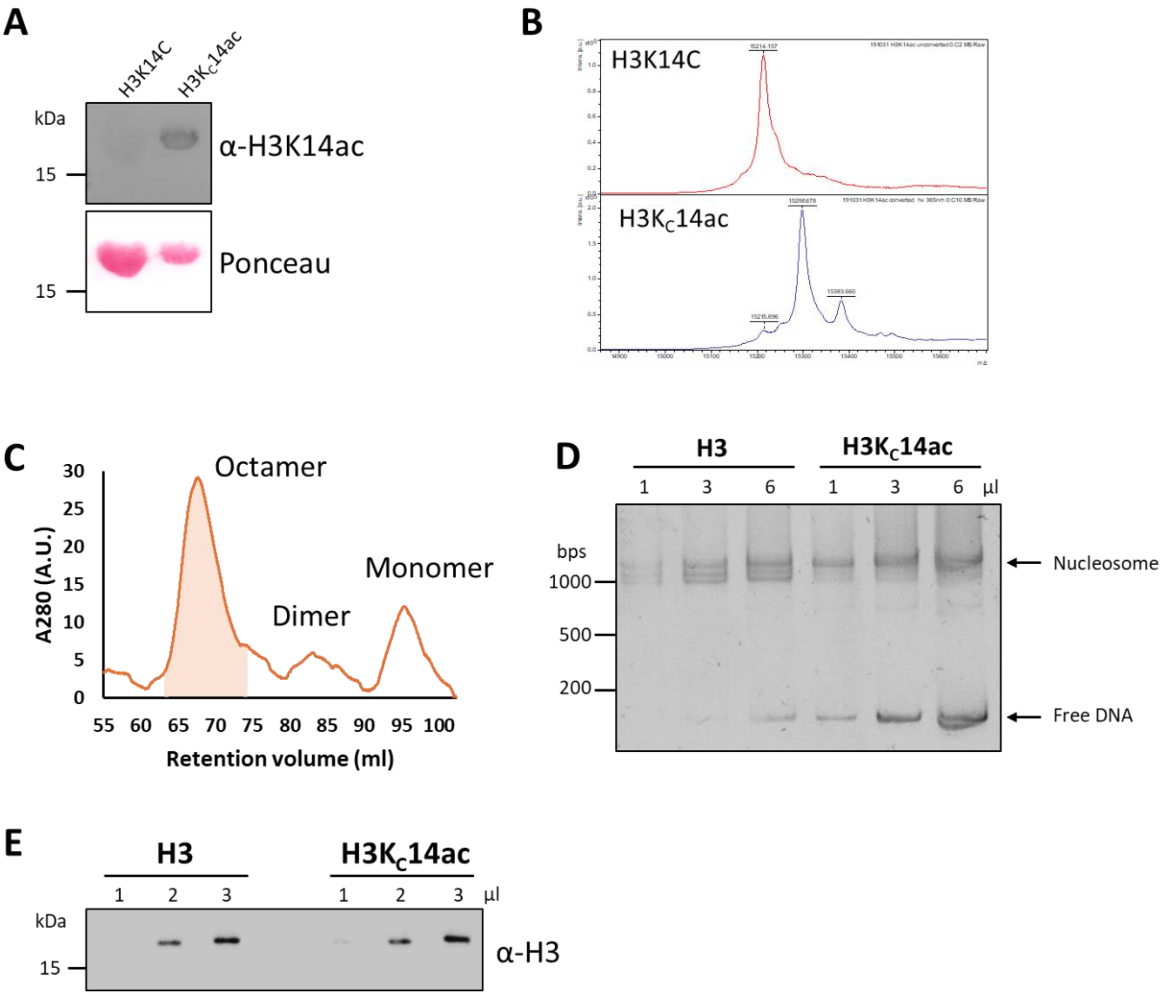


**Supplementary Figure 3: Purification and reconstitution of recombinant nucleosomes.** **A)** Validation of the installation of acetyl lysine analogue at H3K14 with H3K14ac antibody on purified histones. Ponceau staining is used as a loading control. **B)** Mass spectrum of H3 K14C before and after alkylation measured by MALDI-TOF. **C)** Chromatogram of the histone octamer purification by size exclusion chromatography. The fractions that were collected and used for nucleosome reconstitution are highlighted. **D)** Electrophoretic mobility shift assay of reconstituted nucleosome showing the incorporation of DNA into the nucleosomes and free DNA. **E)** Normalization of unmodified and modified nucleosomes by western blot using an H3 antibody.

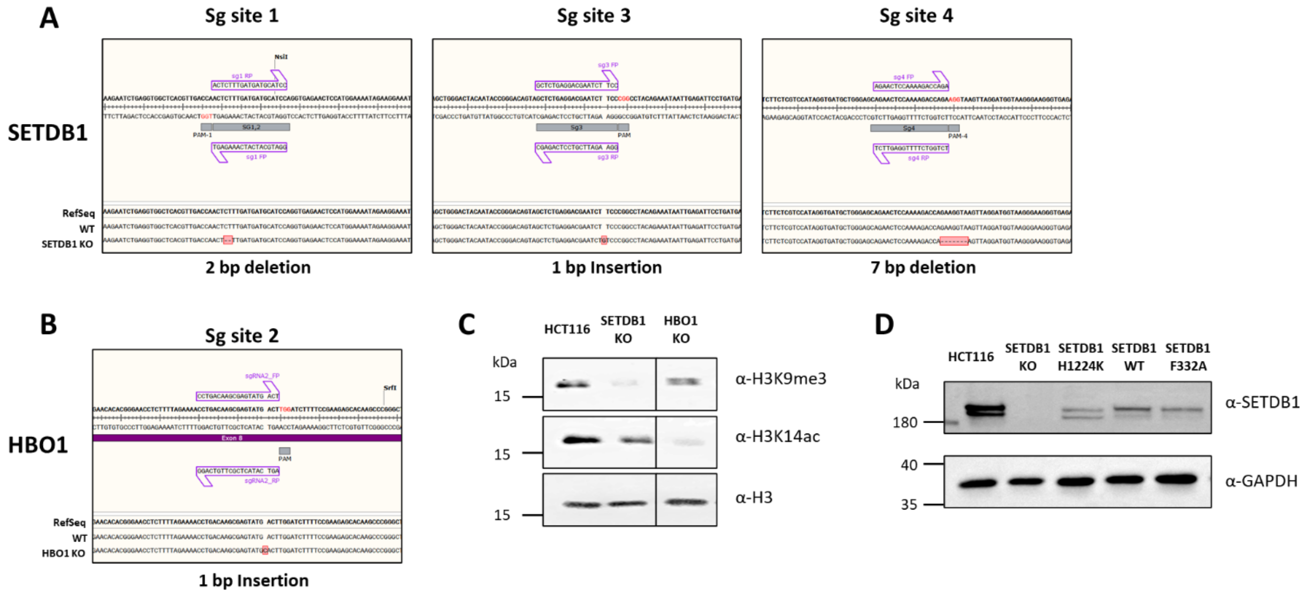


**Supplementary Figure 4: Validation of SETDB1 and HBO1 KO cell clones in HCT116 cells. A)** Sanger sequencing of the PCR product of sgRNA target sites in the SETDB1 KO clone showing a 2 bp deletion at site 1, followed by a 1 bp insertion and 7 bp deletion at site 3 and 4, respectively, all together causing a frameshift in the SETDB1 gene. **B)** Results of corresponding Sanger sequencing of the HBO1 KO clone showing a 1 bp insertion causing a frameshift mutation. **C)** Western blot validation of SETDB1 and HBO1 KO using antibody against H3K9me3 and H3K14ac and H3 used as a loading control. **D)** Western blot validation of SETDB1 rescue experiments with catalytic inactive mutant H1224K, WT and 3TD mutant F332A using SETDB1 antibody. GAPDH is used as a loading control.

**A**

H3K9me3 ChIP-seq

| | | HCT116 | | |
|---|---|---|---|---|
| | | R1 | R2 | Input |
| HCT116 | R1 | 1.00 | 0.98 | 0.68 |
| | R2 | 0.98 | 1.00 | 0.69 |
| | Input | 0.68 | 0.69 | 1.00 |

| | | SETDB1 WT | | |
|---|---|---|---|---|
| | | R1 | R2 | Input |
| SETDB1 WT | R1 | 1.00 | 0.97 | 0.59 |
| | R2 | 0.97 | 1.00 | 0.60 |
| | Input | 0.59 | 0.60 | 1.00 |

| | | SETDB1 KO | | |
|---|---|---|---|---|
| | | R1 | R2 | Input |
| SETDB1 KO | R1 | 1.00 | 0.99 | 0.66 |
| | R2 | 0.99 | 1.00 | 0.64 |
| | Input | 0.66 | 0.64 | 1.00 |

| | | SETDB1 F332A | | |
|---|---|---|---|---|
| | | R1 | R2 | Input |
| SETDB1 F332A | R1 | 1.00 | 0.98 | 0.67 |
| | R2 | 0.98 | 1.00 | 0.68 |
| | Input | 0.67 | 0.68 | 1.00 |

| | | SETDB1 H1224K | | |
|---|---|---|---|---|
| | | R1 | R2 | Input |
| SETDB1 H1224K | R1 | 1.00 | 0.98 | 0.60 |
| | R2 | 0.98 | 1.00 | 0.63 |
| | Input | 0.60 | 0.63 | 1.00 |

| | | HBO1 KO | | |
|---|---|---|---|---|
| | | R1 | R2 | Input |
| HBO1 KO | R1 | 1.00 | 0.94 | 0.62 |
| | R2 | 0.94 | 1.00 | 0.66 |
| | Input | 0.62 | 0.66 | 1.00 |

H3K14ac ChIP-seq

| | | HCT116 | | |
|---|---|---|---|---|
| | | R1 | R2 | Input |
| HCT116 | R1 | 1.00 | 0.84 | 0.55 |
| | R2 | 0.84 | 1.00 | 0.57 |
| | Input | 0.55 | 0.57 | 1.00 |

| | | SETDB1 KO | | |
|---|---|---|---|---|
| | | R1 | R2 | Input |
| SETDB1 KO | R1 | 1.00 | 0.85 | 0.48 |
| | R2 | 0.85 | 1.00 | 0.49 |
| | Input | 0.48 | 0.49 | 1.00 |

**B**

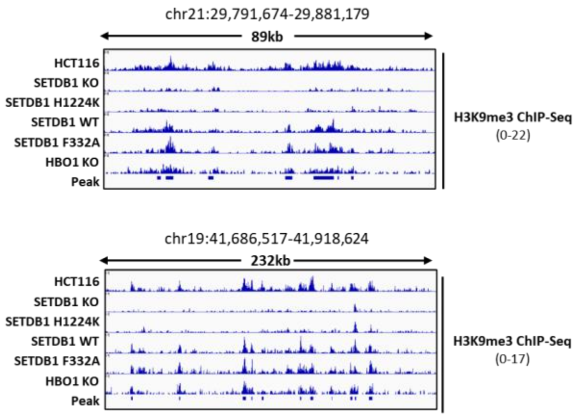


**Supplementary Figure 5: Quality check NGS data analysis. A)** Pearson correlation coefficients showing the reproducibility of H3K9me3 and H3K14ac ChIP-seq experiments between the biological replicates of the indicated cell lines calculated using MultiBigwigSummary of 5 kb bins. **B)** Quality of MACS2 peak calling for the broad H3K9me3 ChIP-seq tracks and merge of all H3K9me3 peaks are shown.

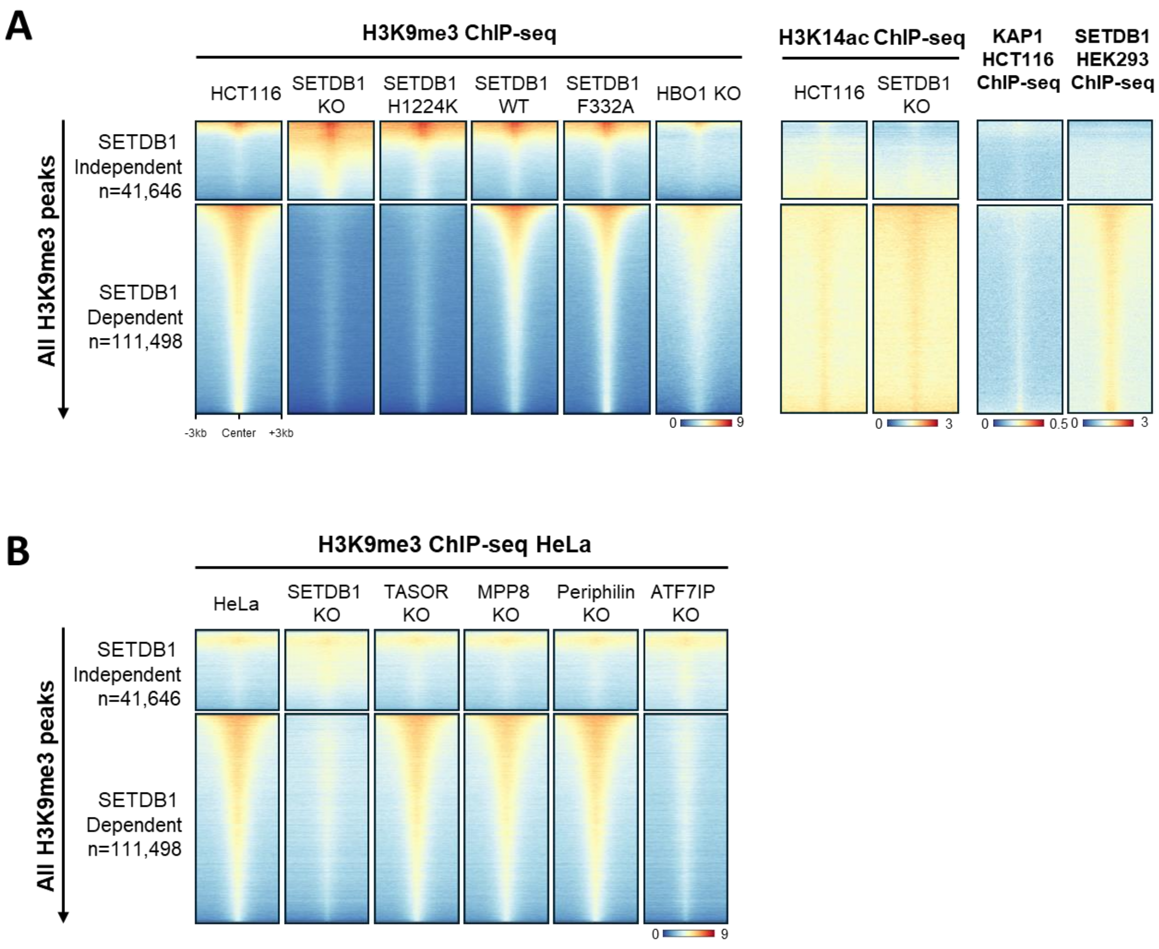


**Supplementary Figure 6: Additional ChIP-seq analysis. A)** ChIP-seq heatmaps related to Figure 4A shown for all H3K9me3 peaks. The number of H3K9me3 peaks in the SETDB1-independent and SETDB1-dependent regions are indicated. ChIP-seq data for KAP1 in HCT116 cells were taken from [1] and ChIP-seq for SETDB1 in HEK293T cells was taken from [2]. **B)** Literature dataset of H3K9me3 ChIP-seq from HeLa cells with KO of HUSH complex subunits [3] or ATFIP [4] clustered based on our differential clustering similar to panel A.

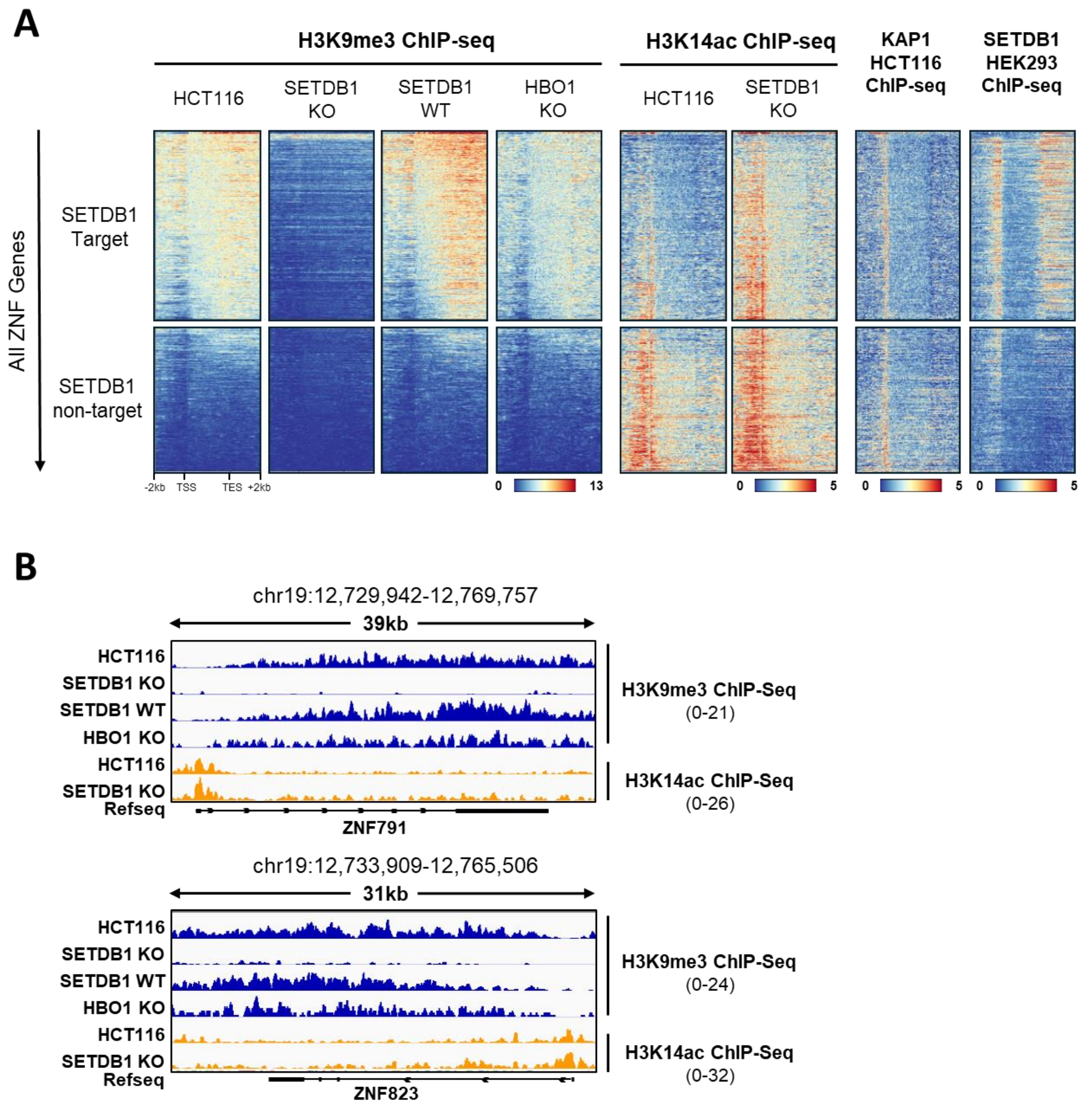


**Supplementary Figure 7: H3K9me3, H3K14ac, KAP1 and SETDB1 ChIP-seq in different cell lines. A)** Heatmap of H3K9me3, H3K14ac and transcription factor ChIP-seq including KAP1 and SETDB1 of different cell lines. Zinc-finger genes were used as regions to plot heatmaps with 2 kb flanking the genes. The previously established clustering of differential H3K9me3 signals into SETDB1 dependent and independent groups in HCT116 cells was used to plot the tracks. **B)** Example IGV browser view images of ChIP-seq profiles at ZNF genes.

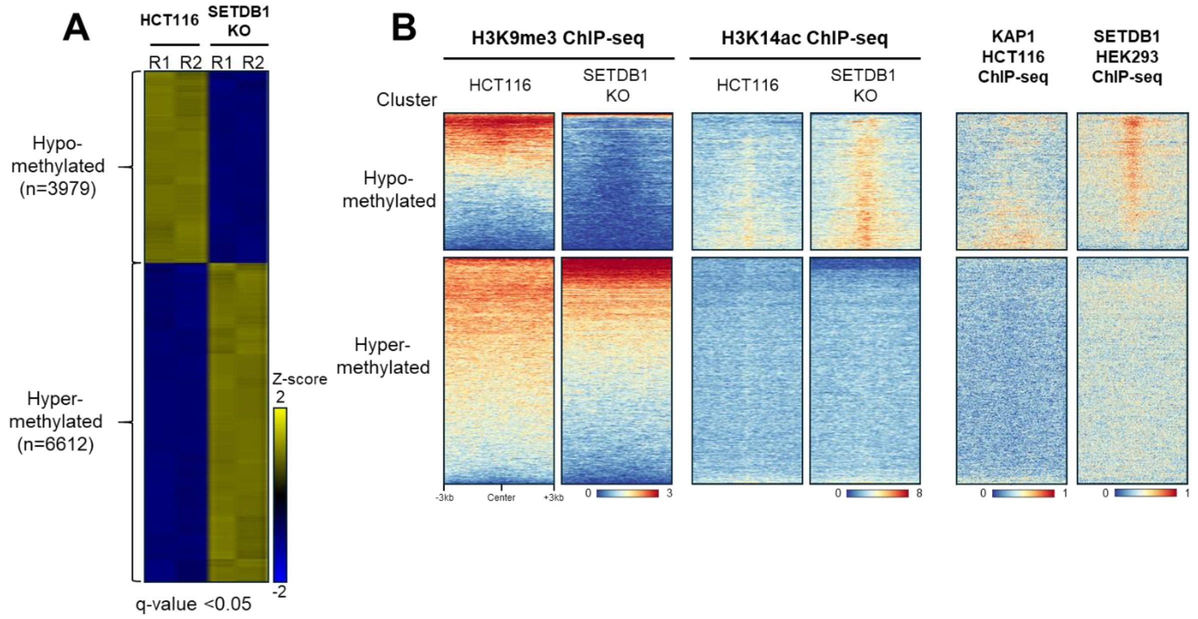


**Supplementary Figure 8: SETDB1 dependent changes in DNA methylation. A)** Differential DNA methylation in HCT116 cells upon SETDB1 KO and the number of CpG is indicated in brackets. T-test statistics were applied to DNA methylation data obtained by Infinium EPIC array analysis in biological replicates using the Qlucore Omics explorer. Statistically significant CpG loci of the biological duplicates (q-value: <0.05, variance >0.4) were used for hierarchical clustering. DNA methylation is presented as z-score. **B)** Hypo- or hypermethylated CpGs in SETDB1 KO (with 3 kb flanking sequence on either side) were used as regions to plot the H3K9me3 and H3K14ac signals along with the occupancy of KAP1 and SETDB1.

### Supplementary Tables

**Supplementary Table 1:** List of peptides used in this study.

| Peptide | H3.1 (aa) | Sequence | Manufacturer |
|---|---|---|---|
| **H3** | 1-18 | $NH_2$-A R T K Q T A R K S T G G K A P R K-$CONH_2$ | Synpeptide |
| **H3K9me2** | 4-19 | Ac-K Q T A R K(me2) S T G G K(Ac) A P R K Q K (Fluorescein) | PEPperPRINT |
| **H3K9me3** | 1-18 | $NH_2$-A R T K Q T A R K(me3) S T G G K A P R K-$CONH_2$ | Synpeptide |
| **H3K14ac** | 1-18 | $NH_2$-A R T K Q T A R K S T G G K(Ac) A P R K-$CONH_2$ | Synpeptide |
| **H3K9me2/ K14ac** | 4-19 | Ac-K Q T A R K(me2) S T G G K(Ac) A P R K Q-K(Fluorescein) | PEPperPRINT |
| **H3K9me3/ K14ac** | 1-18 | $NH_2$-A R T K Q T A R K(me3) S T G G K(Ac) A P R K-$CONH_2$ | Synpeptide |

**Supplementary Table 2:** List of sgRNA used for CRISPR/Cas9 mediated knockout in this study.

| Name | Sequence |
|---|---|
| **sg1-SETDB1_FP** | GGGATGCATCATCAAAGAGT |
| **sg1-SETDB1_RP** | ACTCTTTGATGATGCATCCC |
| **sg2-SETDB1_FP** | GACTCTTTGATGATGCATCC |
| **sg2-SETDB1_RP** | GGATGCATCATCAAAGAGTC |
| **sg3-SETDB1_FP** | GCTCTGAGGACGAATCTTCC |
| **sg3-SETDB1_RP** | GGAAGATTCGTCCTCAGAGC |
| **sg4-SETDB1_FP** | GAGAACTCCAAAAGACCAGA |
| **sg4-SETDB1_RP** | TCTGGTCTTTTGGAGTTCTC |
| **sg1-HBO1_FP** | GGGTGACTCGAGCAGATCGT |
| **sg1-HBO1_RP** | ACGATCTGCTCGAGTCACCC |
| **sg2-HBO1_FP** | CCTGACAAGCGAGTATGACT |
| **sg2-HBO1_RP** | AGTCATACTCGCTTGTCAGG |

**Supplementary Table 3:** List of ChIP-qPCR primers and amplicons used in this study.

| Name | Sequence | Amplicon location (T2T-CHM13v2.0) |
|---|---|---|
| **ERV3-L_FP** | ACCCACACTACGTGGTCGG | chr7:66198993-66199114 |
| **ERV3-L_RP** | TCACAGACTGAGTAGGTTGTC | |
| **CDKL3_FP** | CCACAAACTAACACATCATAC | chr5:134852494-134852593 |
| **CDKL3_RP** | AGTCTGTTGTTAATCTTACCA | |
| **ZNF221_FP** | GGGAAATGTAGTCCAGACGCT | chr19:46774153-46774293 |
| **ZNF221_RP** | TCGTTGAGGGTTCCGAGAGT | |
